# Aiolos restricts the generation of antigen-inexperienced, virtual memory CD8^+^ T cells

**DOI:** 10.1101/2025.06.11.659122

**Authors:** Srijana Pokhrel, Devin M. Jones, Melissa R. Leonard, Jasmine A. Tuazon, Kaitlin A. Read, Gayathri Dileepan, Robert T. Warren, Qiaoke Gong, Jacob S. Yount, Gang Xin, Adriana Forero, Hazem E. Ghoneim, Patrick L. Collins, Emily A. Hemann, Kenneth J. Oestreich

## Abstract

CD8^+^ virtual memory T (T_VM_) cells are memory-like cells that rapidly respond to infection via antigen-independent bystander effector functions. While it is recognized that T_VM_ cells arise independently of foreign antigen encounter, the mechanisms governing their development are not fully understood. Here, we identify the transcription factor Aiolos as a negative regulator of T_VM_ cells. We observed higher quantities of T_VM_ cells in the spleens of uninfected Aiolos-deficient (*Ikzf3^-/-^*) mice relative to wild-type (WT). Furthermore, *Ikzf3^-/-^* T_VM_ cells produced higher levels of IFN-γ and granzyme B. In addition, we found that *Ikzf3^-/-^* T_VM_ cells accumulated to higher quantities in the lungs within 24 hours of influenza virus infection. In line with enhanced T_VM_ functional capacity and lung trafficking, Aiolos-deficient mice cleared virus more rapidly and exhibited reduced morbidity relative to WT animals. Mechanistically, we observed that Aiolos represses the T_VM_ transcriptional regulator Eomes and the IL-15R subunit CD122 (IL-15Rβ/IL-2Rβ), known contributors of T_VM_ cell generation. Collectively, these findings establish Aiolos as a novel molecular repressor of T_VM_ generation and function.

## Introduction

CD8^+^ memory T cells represent a population of immune cells that provide long-term protection from infection^1,2^. These cells are initially generated following antigen encounter during naturally occurring infection or vaccine-induced immune responses and provide the host protection upon reinfection with the same pathogen^1,3,4^. Memory T cells are classified into different subsets, including effector memory (T_EM_), central memory (T_CM_), and tissue-resident memory (T_RM_) cells, each with a distinct niche and set of functions^1,5,6^. While each of these subsets are antigen-experienced, a more recently described subset, termed virtual memory T (T_VM_) cells arise independent of antigen experience^7–9^. Because both T_VM_ and T_CM_ cells express surface markers including L-selectin (CD62L) and the activation marker CD44, T_VM_ cells were previously categorized as a part of the T_CM_ repertoire. However, recent findings have identified that low surface expression of the integrin receptor CD49d, an indicator of foreign antigen-encounter, distinguishes T_VM_ (CD62L^+^CD44^+^CD49d^Lo^) from these other T cell populations^10,11^.

As the generation of T_VM_ population is not dependent upon foreign antigen exposure, they are present in uninfected and unprimed hosts^8^. T_VM_ development and function is dependent upon signals from the cytokine IL-15, as IL-15-deficient mice (*IL-15^-/-^*) lack T_VM_ cells^7,12^. The response to IL-15 in T_VM_ cells is mediated by signals received through the shared receptor subunit CD122 (IL-15Rβ/IL-2Rβ)^7^. Expression of CD122 is positively regulated by the T-box transcription factor, Eomesodermin (Eomes), which is highly expressed in T_VM_ cells and is crucial for their differentiation and survival^7,13^. Eomes binds to the promoter region of *Il2rb* and leads to its activation^14^. While Eomes promotes CD122 expression, IL-15 signaling augments Eomes expression in CD8^+^ T cells^12,15^. Like IL-15 deficiency, Eomes deficiency also results in an absence of T_VM_ cells^13^. Hence, there is an interplay between the key T_VM_ regulators wherein IL-15 signaling upregulates Eomes expression, which in turn promotes CD122 expression in T_VM_ cells, subsequently increasing IL-15 responsiveness.

During infection, T_VM_ cells infiltrate afflicted sites and provide bystander protection independent of TCR engagement^7,10,12,16^. Thus, these cells participate in the first line of defense against various pathogenic infections^17–19^. The protective bystander function of T_VM_ cells depends on the production of inflammatory cytokines (e.g., IFN-γ) and cytotoxic molecules (e.g., granzyme B) in response to cytokines (IL-12 and IL-18), as well as NKG2D-mediated cytotoxicity^7,10^. In addition to their bystander potential, T_VM_ cells can also differentiate into early effector and late memory T cells in response to specific antigens, as has been previously demonstrated in the case of influenza virus infection^17^. Their polyclonal repertoire allows for antigen-specific clonal expansion, and they are poised to respond to an antigen more efficiently than naive CD8^+^ T cells^7,11,17,20^. T_VM_ cells have also been implicated in the resolution of cancer, as these cells can infiltrate tumors and target cancer cells independent of MHC-I mediated recruitment and response ^13,21^. Also, despite having a high avidity for self-antigen, which could be a concern for autoimmunity, these cells demonstrate high self-tolerance and promote homeostasis^22^. Together, these rapid, antigen-independent responses, coupled with the ability to differentiate in an antigen-specific manner, make T_VM_ cells a unique therapeutic target for the treatment of a range of diseases. Hence, obtaining a better understanding of the full repertoire of transcriptional networks that regulate their development and function is beneficial.

Complex transcriptional networks regulate the generation of different memory subsets^23–25^. We have recently shown that the Ikaros Zinc Finger (IkZF) transcription factor Aiolos (encoded by *Ikzf3*) represses Eomes expression and hinders cytotoxic potential in CD4^+^ cytotoxic T lymphocytes (CTLs), yet promotes CD4^+^ T follicular helper (T_FH_) cell transcriptional programming^26,27^. While our previous work and that of others have established the contribution of different IkZF factors to CD4^+^ T cell differentiation, knowledge regarding their role in CD8^+^ T cell differentiation is limited ^26–32^. Given that similar cytokine signaling pathways and transcription factors regulate the differentiation and function of these two cell types, it is relevant and important to investigate how IkZF factors impact the differentiation of CD8^+^ T cells^33,34^.

Here, we identify Aiolos as a negative regulator of Eomes, CD122 expression, and CD8^+^ T_VM_ cell generation. We observed significantly higher numbers of T_VM_ cells in the spleens of uninfected Aiolos-deficient mice compared to WT. This increased number of T_VM_ cells at baseline subsequently translated into a significant increase in the resident T_VM_ cell population in the lungs of Aiolos-deficient mice 1 day post influenza virus infection. Phenotypically, Aiolos-deficient T_VM_ cells displayed increased Eomes and CD122 expression, as well as enhanced production of IFN-γ and granzyme B. Using a murine model of influenza virus infection, we found that T_VM_ cells contributed to reduced disease severity in Aiolos-deficient mice compared to WT animals, which manifested as lower weight loss and viral titers at early timepoints post-infection. Together, these findings support a previously unappreciated role for Aiolos in dampening CD8^+^ T_VM_ responses during viral infection.

## Results

### Aiolos suppresses Eomes and CD122 expression in CD8^+^ T cells

We previously established that loss of Aiolos in effector CD4^+^ T cells during influenza A virus infection (A/PR/8/34 (H1N1), termed PR8) results in augmented expression of the transcription factor Eomes and induction of the CD4-CTL gene program^26^. Given the prominent role of Eomes in CD8^+^ T cell biology, including its ability to induce both cytotoxic features and memory cell programming, we sought to determine whether this relationship between Aiolos and Eomes may be a conserved feature in CD8^+^ T cells^14,35,36^. Initially, we examined Eomes protein expression in CD8^+^ T cells isolated from the spleens of uninfected, specific-pathogen free (SPF) wild type (WT) and Aiolos-deficient (*Ikzf3^-/-^)* C57BL/6J mice using flow cytometry. In the absence of Aiolos, both the percentage of Eomes^+^ T cells, and Eomes protein expression were significantly higher in Aiolos-deficient CD8^+^ T cells compared to WT controls (Fig. 1a-c). Further analysis of CD8^+^ T cells revealed increased percentages of Eomes^+^ naive (CD62L^+^CD44^-^) and activated (CD44^+^) T cells in Aiolos-deficient mice compared to WT (Supplementary Fig. 1a-d). In addition, Aiolos deficiency also impacted the expression of IL-15R subunit CD122, an Eomes target gene encoded by *Il2rb*^14^. We observed an increase in the percentage and numbers of CD122^+^ T cells in Aiolos-deficient animals compared to WT controls (Fig. 1d-e). To further define the role of Aiolos in regulating Eomes and CD122 expression, we overexpressed Aiolos in *Ikzf3^-/-^*CD8^+^ T cells *in vitro* using retroviral transduction (Supplementary Fig. 2a-b). Overexpression of Aiolos resulted in a significant decrease in the expression of Eomes as well as CD122 (Supplementary Fig. 2c-d). Collectively, these *in vivo* and *in vitro* data demonstrate that Aiolos represses Eomes and CD122 expression in primary CD8^+^ T cells in a cell-intrinsic manner.

**Figure 1.**
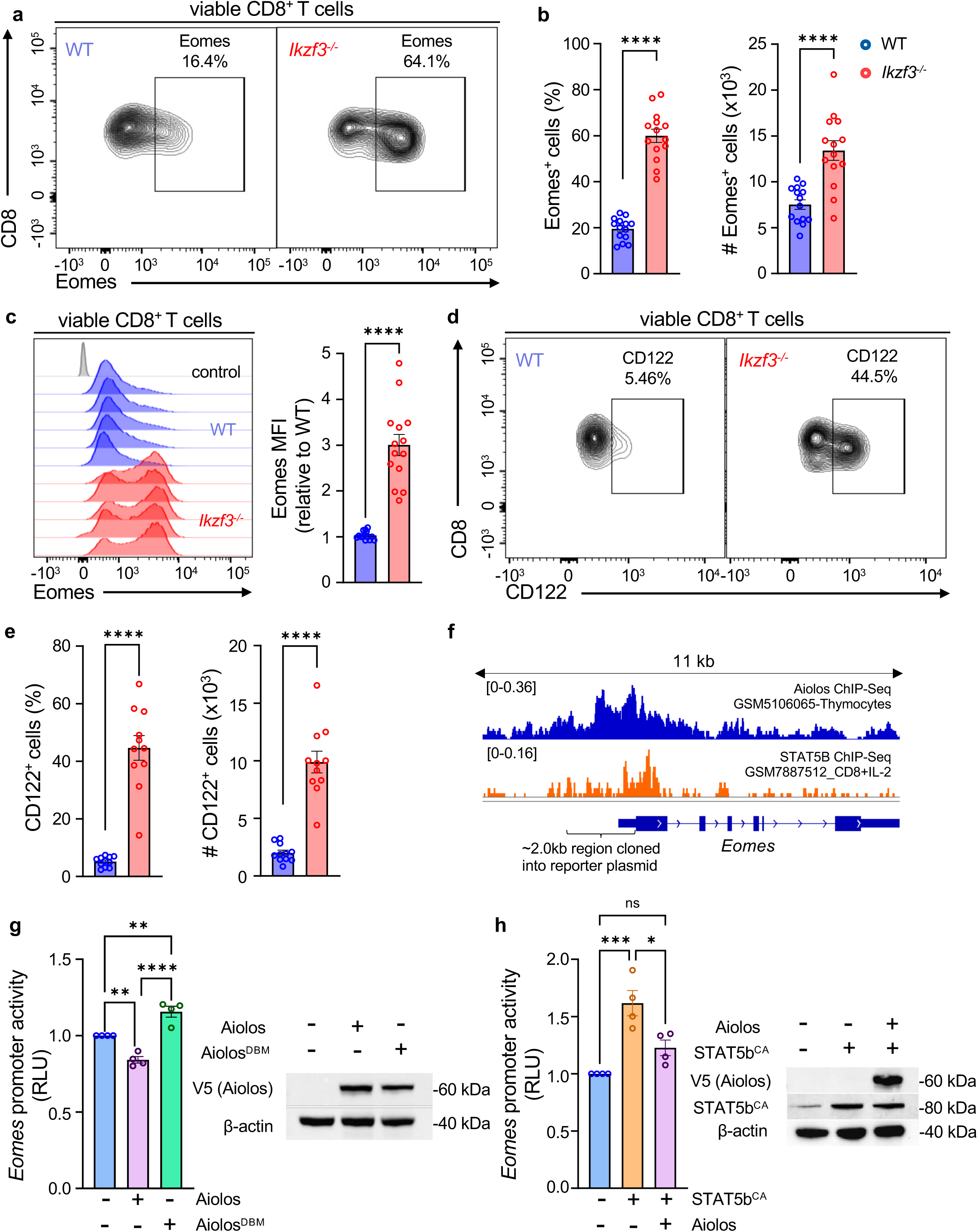
Aiolos expression inversely correlates with that of Eomes and CD122 in CD8^+^ T cells. Analysis of Eomes expression in CD8^+^ T cells isolated from the spleens of uninfected WT and *Ikzf3^-/-^* mice. (**a**) Representative flow contour plots depicting expression of Eomes in WT and *Ikzf3^-/-^* CD8^+^ T cells; and (**b**) respective bar graphs showing percent (%) of Eomes^+^ and numbers of Eomes-expressing CD8^+^ T cells (#: counts, normalized to 6 x 10^5^ events). (**c**) Representative histogram overlay for Eomes expression and associated data showing differences in median fluorescence intensity (MFI) fold change relative to WT control. (**a-c**) Data shown for 4 independent experiments, n=14, mean ± SEM, unpaired Student’s t-test, ****p≤0.0001. (**d**) Representative flow plots (**e**) associated bar graphs showing percent (%) of CD122^+^ and numbers of CD122-expressing CD8^+^ T cells (#: counts, normalized to 6 x 10^5^ events), Data shown for 2 independent experiments, n=8, mean ± SEM, unpaired Student’s t-test, ****p≤0.0001. (**f**) Analysis of publicly available Aiolos Chromatin Immunoprecipitation (ChIP)-Seq data (GSM5106065) and STAT5b ChIP-Seq data (GSM7887512) showing enrichment of Aiolos and STAT5b at the *Eomes* promoter region. Sequencing tracks were viewed using Integrative Genomics Viewer. The gene region cloned into a reporter vector for Eomes promoter activity is indicated. (**g-h**) EL4 T cells were transfected with an *Eomes* promoter-reporter construct in conjunction with vectors for Aiolos, Aiolos^DBM^, STAT5b^CA^, or empty vector control. As a control for transfection efficiency, cells were concurrently transfected with SV40-*Renilla*, and luciferase activity was used as a readout for promoter activity. Luciferase promoter-reporter values were normalized to SV40-*Renilla* control and presented as relative to empty vector. Immunoblot analysis of indicated proteins confirming the overexpression of respective vectors. β-actin was used as a loading control. Data are representative of 4 independent experiments, n=4, mean ± SEM, one-way ANOVA with Tukey’s multiple comparisons test, *p≤0.05, **p≤0.01, ***p≤0.001, and ****p≤0.0001.

To determine the mechanism by which Aiolos suppresses Eomes, we analyzed publicly available Aiolos ChIP-Seq data from murine thymocytes [GSM5106065^37^]. Analysis of Aiolos enrichment across the *Eomes* gene using Integrative Genomics Viewer (IGV) revealed peaks at the *Eomes* promoter region, suggesting that Aiolos may directly repress its expression (Fig. 1f). To assess the potential impact of Aiolos on *Eomes* promoter activity, we generated plasmids to overexpress Aiolos or an Aiolos DNA-binding mutant (Aiolos^DBM^), along with a firefly luciferase gene reporter construct containing the *Eomes* promoter region. Aiolos overexpression in EL4 T cells resulted in suppression of *Eomes* promoter activity (Fig. 1g). However, the repressive effect was lost with Aiolos^DBM^ which lacks a functional DNA binding domain (Fig. 1g), suggesting that direct DNA binding is necessary for the suppressive effect of Aiolos. A known driver of Eomes expression in CD8^+^ T cells is the transcription factor STAT5^38^. Interestingly, sites of STAT5b enrichment [GSM7887512^39^] at the Eomes promoter region overlapped with the Aiolos enrichment sites (Fig. 1f). This is in line with previously published studies, including work from our lab, indicating that IkZF and STAT factor DNA-binding motifs overlap^26,40,41^. To determine whether Aiolos may repress Eomes expression by interfering with STAT5 mediated induction of Eomes, we overexpressed a constitutively active form of STAT5b (STAT5b^CA^) along with Aiolos in EL4 T cells. As expected, STAT5b overexpression alone significantly enhanced *Eomes* promoter activity. However, co-expression with Aiolos abrogated the STAT5-induced response (Fig. 1h).

Together, these findings support Aiolos as a transcriptional repressor of Eomes in CD8^+^ T cells through direct interaction or via competition with STAT5 for promoter occupancy. Further, our data also suggests the suppression of CD122 by Aiolos in these cells which could result in their decreased cytokine responsiveness. Given the interplay between IL-15 signaling and Eomes, these findings are suggestive of a mechanism whereby Aiolos regulates the IL-15/STAT5/Eomes axis.

### Inverse correlation of Aiolos versus Eomes and CD122 expression within CD8^+^ T_VM_ cells

Both Eomes and CD122 are important for CD8^+^ T_VM_ cell generation^7,12,13^. To understand the potential interplay between Aiolos, Eomes and CD122 in regulating T_VM_ cells, we began by examining Aiolos, Eomes and CD122 expression in the spleen-resident CD8^+^ T cell subsets in uninfected WT mice: naive (CD62L^+^CD44^-^), effector (T_EFF_, CD62L^-^CD44^+^), and T_VM_ (CD62L^+^CD44^+^ CD49d^Lo^). To distinguish between circulating (CD45.2^(i.v.Pos))^ and non-circulating resident (CD45.2^(i.v. Neg)^) cells, we retro-orbitally injected mice with fluorophore-labeled anti-CD45.2 antibody prior to euthanasia (Supplementary Fig. 3a). Analysis of the resident CD8^+^ T cell populations revealed increased expression of Eomes and CD122 in T_VM_ cells relative to T_EFF_ cells, while Aiolos expression exhibited the opposite trend (Supplementary Fig. 3b-c). The inverse correlation between Aiolos expression and that of both Eomes and CD122 suggests that, similar to our *in vitro* findings, Aiolos may function to negatively regulate these key molecules in T_VM_ populations *in vivo*.

### Aiolos deficiency results in an elevated splenic T_VM_ population in uninfected mice

We next sought to determine the impact of Aiolos deficiency on T_VM_ cell generation. As T_VM_ cells are known to be present in the spleens of unprimed hosts, we assessed the CD8^+^ T_VM_ population in the spleens of uninfected WT and Aiolos-deficient mice^7,9^. We observed a significant increase in both the frequency and numbers of resident CD8^+^ T_VM_ cells in the spleens of Aiolos-deficient mice compared to WT (Fig. 2a-b). This was consistent with our prior observation that Eomes and CD122 were significantly elevated in Aiolos-deficient bulk CD8^+^ T cells (Fig. 1a-e). As noted previously, T_VM_ cells exhibit elevated expression of both Eomes and CD122 relative to the naive CD8^+^ T cells. Yet, we observed a further increase in their expression in Aiolos-deficient T_VM_ cells compared to WT (Fig. 2c-d). Together, these findings support a repressive role for Aiolos in T_VM_ cell differentiation, potentially through modulation of Eomes and CD122.

**Figure 2.**
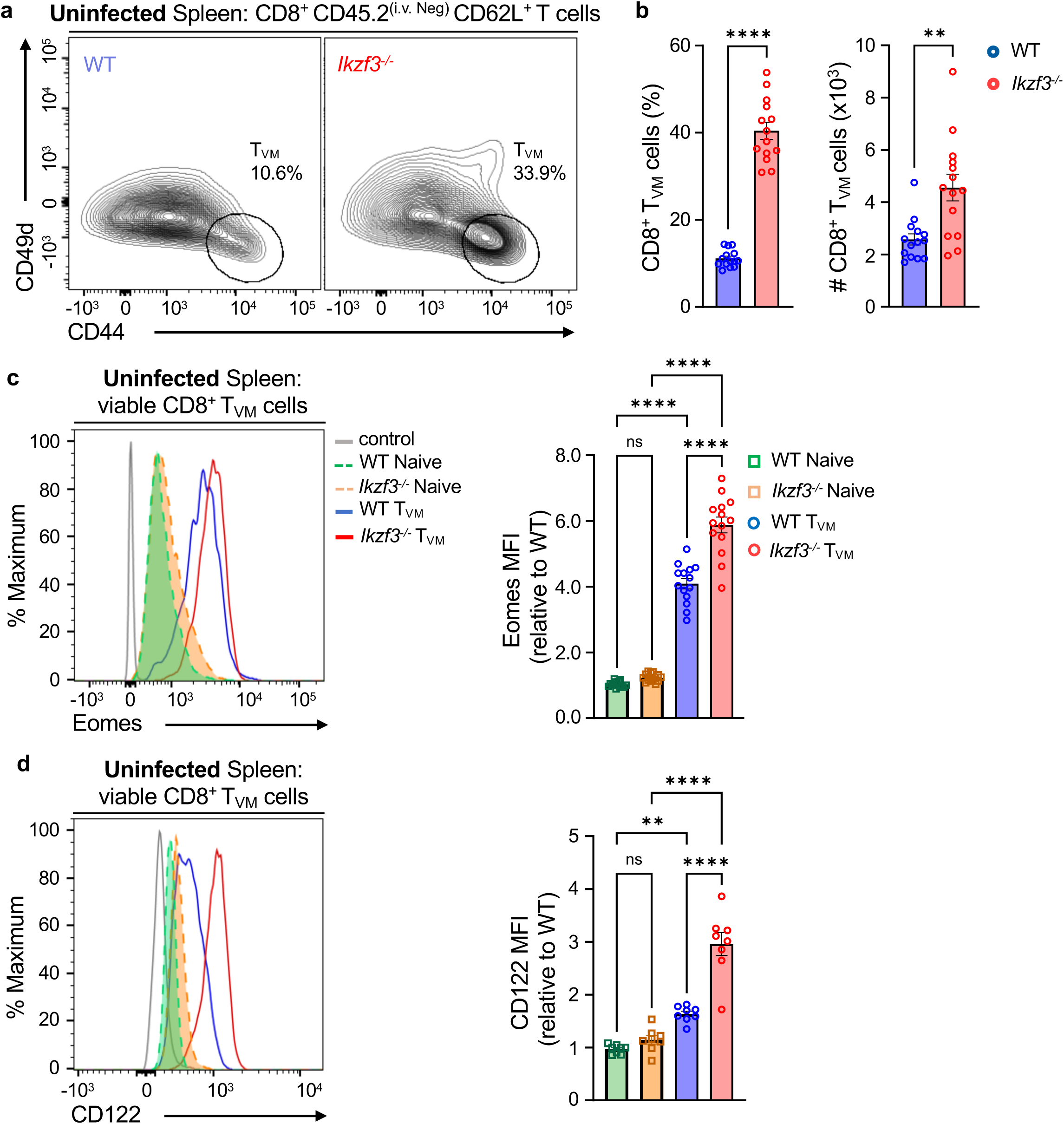
Aiolos deficiency results in elevated splenic T_VM_ populations. Flow cytometry analysis of CD8^+^ T cells from the spleens of uninfected WT and *Ikzf3^-/-^* mice. Mice were injected with fluorophore (PE) labeled anti-CD45.2 antibody via the retro-orbital route to distinguish resident (CD45.2^(i.v. Neg)^) from circulating (CD45.2^(i.v. Pos)^) T_VM_ cells. (**a**) Representative flow contour plots depicting CD8^+^ T_VM_ (CD62L^+^CD44^+^CD49d^Lo^) cells from WT and *Ikzf3^-/-^* mice and (**b**) corresponding data for percent (%) and counts (#, normalized to 6 x 10^5^ events) of CD8^+^ T_VM_ cells. Data shown for 4 independent experiments, n=14, mean ± SEM, unpaired Student’s t-test, **p≤0.01, and ****p≤0.0001. (**c-d**) Representative histogram overlays and corresponding median fluorescence intensity (MFI) fold change for (**c**) Eomes and (**d**) CD122 in the indicated populations, relative to WT naive CD8^+^ T cells. Data shown for 2-4 independent experiments, n=14 (for c) and n=8 (for d), mean ± SEM; two-way ANOVA with Tukey’s multiple comparisons test, **p≤0.01, and ****p≤0.0001.

### Aiolos deficiency promotes the T_VM_ gene program

To determine how Aiolos may regulate the transcriptome of T_VM_ populations, we performed RNA-sequencing analysis on T_VM_ cells isolated from the spleens of uninfected WT and Aiolos-deficient mice. We also sequenced RNA from WT naive CD8^+^ T cells as control. Principal Component Analysis (PCA) of transcripts for T_VM_ cells and WT naive CD8^+^ T cells revealed distinct clusters between the naive and T_VM_ samples, as has been previously reported^7^. Importantly, PCA analysis of the T_VM_ transcripts revealed that Aiolos-deficient T_VM_ samples formed a cluster distinct from WT T_VM_ (Fig. 3a). Hierarchical clustering of the top 200 differentially expressed genes identified unique gene expression patterns in WT and Aiolos-deficient T_VM_ cells (Fig. 3b). In Aiolos-deficient T_VM_ cells, genes involved in T_VM_ transcriptional regulation (*Eomes*, *Tbx21*) were upregulated along with cytokine receptors known to promote T_VM_ homeostasis and responsiveness to IL-15 (*ll15ra, Il2rb)* (Fig. 3c-d)^7,12^. This increase in *Eomes* and *Il2rb* transcript was consistent with our previous findings of elevated Eomes and CD122 protein levels in Aiolos-deficient samples (Fig. 2c-d). In line with these, gene set enrichment analysis (GSEA) of immunologic signature gene sets revealed that effector genes that were downregulated in IL-2Rβ deficient CD8^+^ T cells were upregulated in Aiolos-deficient T_VM_ samples, supporting the inverse relationship between Aiolos and IL-2Rβ (Fig. 3e). In addition, genes involved in T_VM_ cell infiltration into infected tissues (*Ccr2*, *Cxcr3 and S1pr5*) were upregulated in the absence of Aiolos, whereas other chemokine receptor genes such as *Ccr9* and *Cxcr5* were downregulated in Aiolos-deficient T_VM_ samples (Fig. 3c-d)^7,17^.

**Figure 3.**
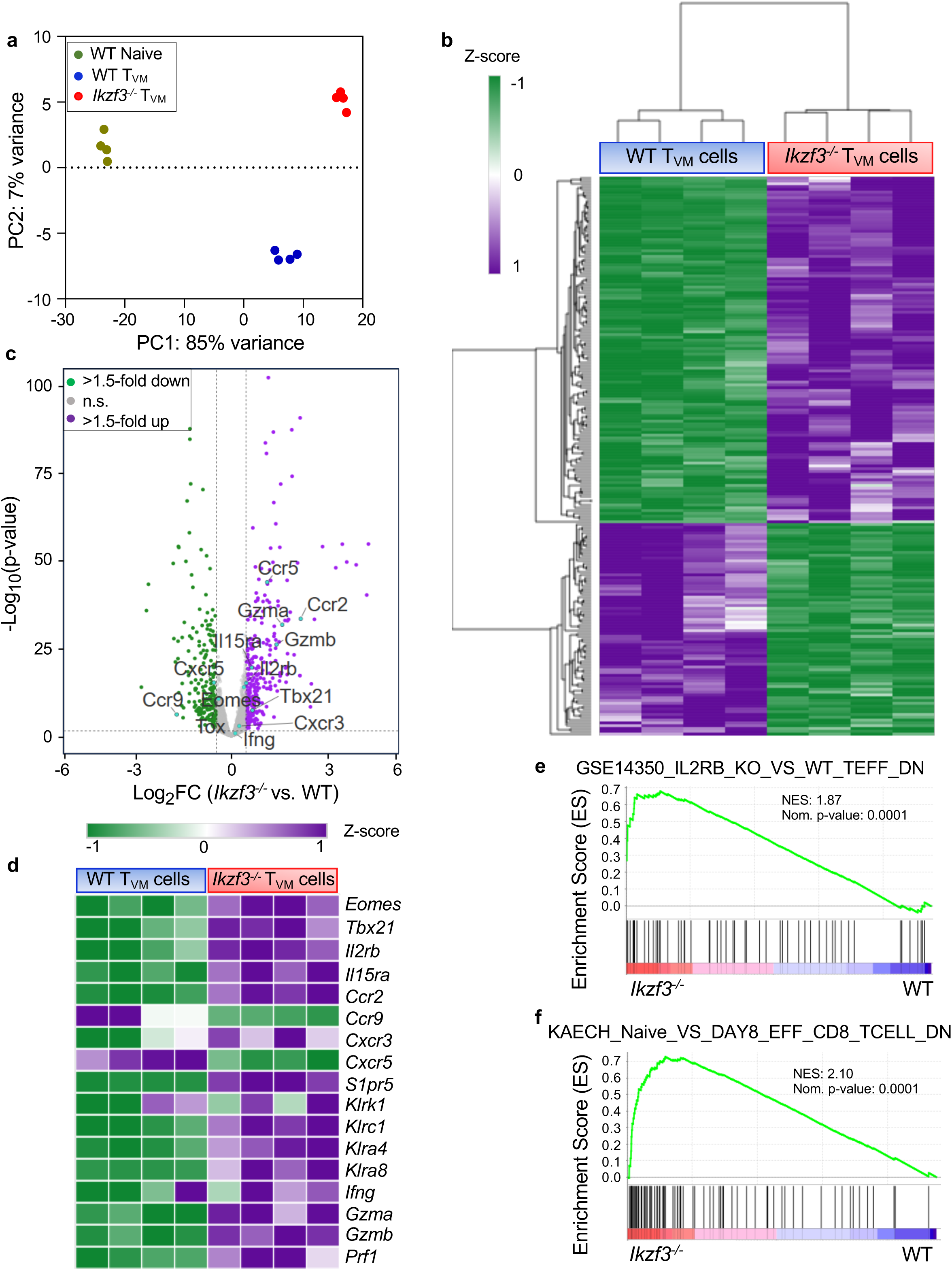
Aiolos deficiency enhances features of the T_VM_ gene program. CD8^+^ T_VM_ cells were sorted from splenic CD8^+^ T cells of WT and *Ikzf3^-/-^* mice. Similarly, naive CD8^+^ T cells were isolated from the spleens of WT mice and RNA-seq analysis was performed to assess differentially expressed genes (DEGs) between WT and Aiolos-deficient cells. (**a**) PCA analysis of normalized DESeq2counts. Data representative of 4 independent experiments. (**b**) Heatmap of top 200 DEGs between WT and Aiolos-deficient T_VM_ cells clustered by Euclidian distance. Changes in gene expression are presented as row (gene) Z-score. (**c**) Volcano plot displaying gene expression changes in WT vs. Aiolos-deficient cells. Genes were color-coded as follows: no significant changes in expression (gray), upregulated genes with >1.5-fold change in expression with a P < 0.05 (purple), downregulated genes with >1.5-fold change in expression with a P < 0.05 (green) and selected genes of interest (turquoise). (**d**) Representative heatmap of DEGs that are positively and negatively associated with the T_VM_ gene program in WT vs. Aiolos-deficient cells. (**e-f**) Gene Set Enrichment Analysis (GSEA) analysis of pre-ranked (sign of fold change X −log10 (p-value)) genes using the Broad Institute GSEA software for comparison against immunological signature gene sets. Enrichment plots for indicated gene sets are shown.

We also analyzed the protein levels of transcription factor T-bet (encoded by *Tbx21*) and the migratory receptor CXCR3 (encoded by *Cxcr3)* in naive CD8^+^ and T_VM_ cells^14,17^. As expected, we observed increased expression of these factors in T_VM_ cells compared to the naive counterparts. Importantly, their expression was significantly higher in Aiolos-deficient T_VM_ cells when compared to WT T_VM_ (Supplementary Fig. 4a-b), corroborating our findings from the RNA-seq analysis.

In addition to increased tissue infiltration, CCR2-expressing T_VM_ cells are reported to exhibit enhanced cytotoxic potential compared to the CCR2-deficient T_VM_ cells^17^. To determine the impact of Aiolos on T_VM_ cytotoxicity, we analyzed the expression of cytotoxic genes expressed in T_VM_ cells, the NK cell genes involved in cell killing (*Klrk1, Klrc1, Klra4, and Klra8*) as well cytotoxic (*Gzma, Gzmb, Prf1*) and proinflammatory (*Ifng*) genes^9,42,43^. Transcript analysis revealed increased expression of these genes in Aiolos-deficient T_VM_ samples compared to their WT counterparts (Fig 3c-d). Similarly, GSEA analysis of gene ontology and immunologic signature subsets also revealed that genes involved in the regulation of innate immune responses as well as effector and memory response were upregulated in Aiolos-deficient T_VM_ cells (Fig. 3f and Supplementary Fig. 4c-d), suggestive of a potential for functional superiority of T_VM_ cells in the absence of Aiolos.

### Aiolos deficiency augments the functional properties of T_VM_ cells

One of the functional characteristics of T_VM_ cells is to provide rapid bystander protection through the production of IFN-γ and granzyme B in response to cytokine stimulation (IL-12, IL-18, and IL-15), independent of T cell receptor activation or co-stimulation^7,10^. In the absence of Aiolos, we observed that genes encoding IL-15R subunits (*Il15ra* and *Il2rb*), IL-12R subunits (*Il12rb1, Il12rb2*), and IL-18R subunits (*Il18r1 and Il18rap*) were upregulated in T_VM_ cells suggesting that Aiolos deficiency may augment T_VM_ responsiveness to these cytokines (Fig. 3c, 4a). GSEA analysis of the immunologic signatures also revealed that genes expressed in the presence of IL2Rβ/IL15Rβ and in IL-12-stimulated effector CD8^+^ T cells were upregulated in Aiolos-deficient T_VM_ cells (Fig. 3e, and Supplementary Fig. 4e). To determine whether this increase in receptor genes resulted in a functional difference in cytokine signaling, we assessed the tyrosine phosphorylation-mediated activation of STAT5 and STAT4, which are involved in IL-15 and IL-12 signaling, respectively. We isolated bulk CD8^+^ T cells from the spleens of uninfected WT and *Ikzf3^-/-^* mice and treated them with a cytokine cocktail (IL-12, IL-18, and IL-15C) *ex vivo* (Fig. 4b). We then determined the phosphorylation status of STAT5 (pY-STAT5) and STAT4 (pY-STAT4) in T_VM_ cells via flow cytometry. We observed elevated STAT5 and STAT4 phosphorylation in the Aiolos-deficient T_VM_ cells compared to WT, suggestive of an enhanced cytokine responsiveness in the absence of Aiolos (Fig. 4c-h).

**Figure 4.**
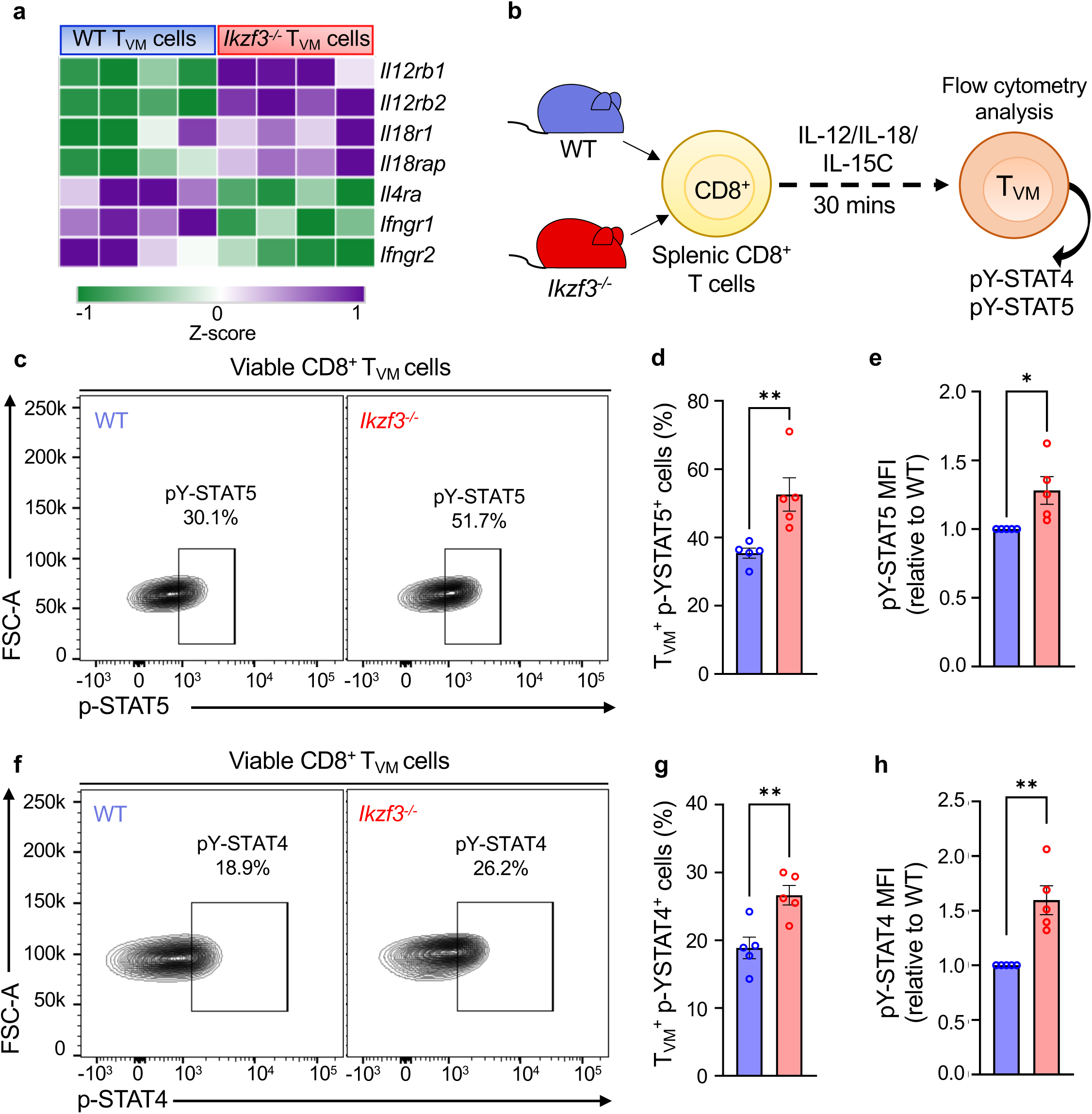
Loss of Aiolos results in enhanced cytokine responsiveness in T_VM_ cells. (**a**) Heatmap showing differences in different cytokine receptors between WT and Aiolos-deficient T_VM_ samples. (**b**) Schematic of the *in vitro* treatment of WT and Aiolos-deficient splenic CD8^+^ T cells. Bulk CD8^+^ T cells were isolated from the spleens of WT and Aiolos-deficient mice and treated with rmIL-12, rmIL-18 and IL-15C (IL-15/IL-15Rα complex) for 30 minutes. Cells were then immediately fixed with formaldehyde and permeabilized with methanol for pY-STAT5 and pY-STAT4 staining. (**c, f**) Representative flow contour plots and (**d, g**) associated data graphs for pY-STAT5 and pY-STAT4 between WT and Aiolos-deficient T_VM_ cells, relative to WT. (**e, h**) Median fluorescence intensity (MFI) fold change for pY-STAT5 and pY-STAT4 between WT and Aiolos-deficient T_VM_ cells, relative to WT. Data representative of 5 independent experiments, n=5, mean ± SEM; unpaired Student’s t-test, *p≤0.05, and **p≤0.01.

Next, to assess whether the enhanced cytokine signaling had a functional consequence, we examined production of IFN-γ and granzyme B by WT and Aiolos-deficient T_VM_ cells. We treated CD8^+^ T cells from these mice with the cytokine mixture as above for 16 hours and then analyzed IFN-γ and granzyme B production by T_VM_ cells via flow cytometry (Fig. 5a). Indeed, Aiolos-deficient T_VM_ cells produced significantly elevated levels of IFN-γ and granzyme B (Fig. 5b-c, and 5e-f) and had higher expression of both factors (Fig. 5d and g) compared to WT T_VM_ cells. Notably, both factors are regulated by T-bet, the expression of which was also elevated in Aiolos-deficient T_VM_ (Supplementary Fig. 4a). Collectively, these data demonstrate that Aiolos deficiency results in increased numbers of T_VM_ cells with greater functional capacity.

**Figure 5.**
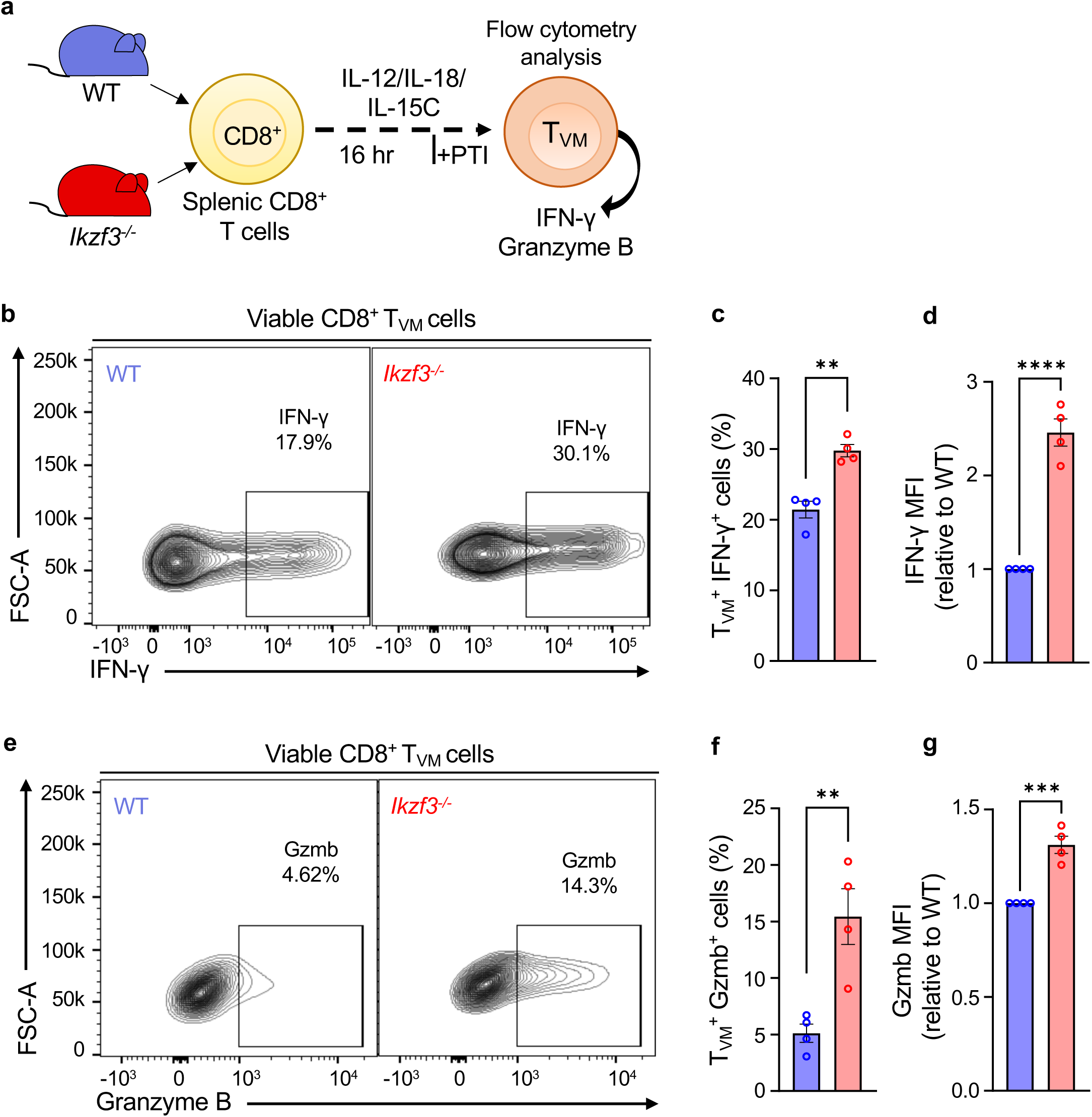
Aiolos deficiency promotes effector functions of T_VM_ cells. (**a**) Schematic of the *in vitro* treatment of WT and Aiolos-deficient splenic CD8^+^ T cells to determine IFN-γ and granzyme B production by CD8^+^ T_VM_ cells. Splenic CD8^+^ T cells were treated with the cytokines (IL-12, IL-18 and IL-15C) for 16 hours. Protein transport inhibitor (PTI) was added 4 hours prior to harvesting the cells for flow cytometry analysis. Representative flow plots depicting (**b**) IFN-γ and (**e**) granzyme B production by WT and Aiolos-deficient T_VM_ cells in response to cytokine stimulation. Data for percent (%) of (**c**) IFN-γ and (**f**) granzyme B production by CD8^+^ T_VM_ cells. Median fluorescence intensity (MFI) fold change for (**d**) IFN-γ and (**g**) granzyme B, relative to WT. Data representative of 4 independent experiments, n=4, mean ± SEM; unpaired Student’s t-test, **p≤0.01, ***p≤0.001 and ****p≤0.0001.

### Aiolos deficiency enhances the response of CD8^+^ T_VM_ cells to influenza virus infection

As a part of their protective response, CD8^+^ T_VM_ cells are known to infiltrate the site of infection and provide bystander protection^7,10,12,16^. In line with this feature, Qi and colleagues have demonstrated that CD8^+^ T_VM_ cells can be found in influenza virus-infected lungs as early as 1 day post-infection (1 d.p.i.)^17^. To determine whether the absence of Aiolos would augment this response, we infected WT and Aiolos-deficient mice with the mouse-adapted H1N1 influenza A virus strain PR8 and assessed T_VM_ populations at 1 d.p.i. Similar to our observations in uninfected mice, the frequency and numbers of resident T_VM_ cells were increased in the spleens of Aiolos-deficient mice 1 d.p.i. compared to WT (Fig. 6a-b). Strikingly, there was also a significant increase in the number of resident CD8^+^ T cells in the lungs of Aiolos-deficient mice compared to WT (Fig. 6c-d). Further phenotypic analyses of these populations for naive, effector and T_VM_ cells revealed that T_VM_ cells represented the majority of resident CD8^+^ T cells in both WT and Aiolos-deficient animals (Fig. 6e). Consistent with our previous findings, Aiolos-deficient T_VM_ cells from infected spleens and lungs exhibited elevated expression of both Eomes and CD122 at 1 d.p.i. compared to WT (Supplementary Fig. 5a-b).

**Figure 6.**
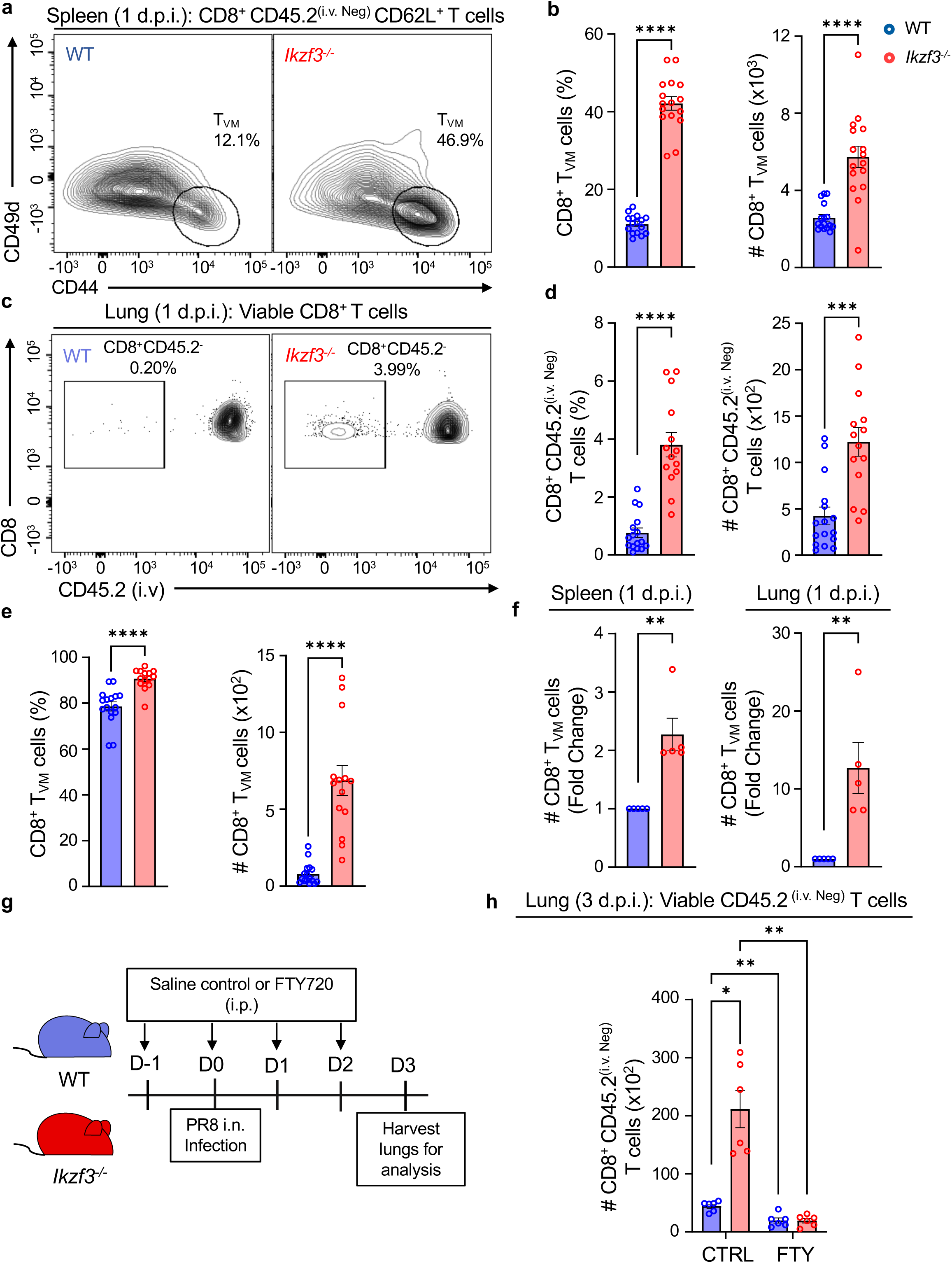
Aiolos suppresses the response of CD8^+^ T_VM_ cells to influenza A virus infection. WT and Aiolos-deficient mice were intranasally (i.n.) infected with 100 plaque-forming units (PFU) of influenza A virus (A/PR/8/34, or “PR8”) and tissues were harvested 1 day post-infection (d.p.i). (**a**) Representative flow contour plot depicting resident (CD45.2^(i.v. Neg)^) CD8^+^ T_VM_ cells in the spleens of WT and *Ikzf3^-/-^* mice 1 d.p.i. and (**b**) corresponding percent (%) positive and counts (#, normalized to 6 x 10^5^ events) data shown for CD8^+^ T_VM_ cells. (**c**) Representative flow plots and (**d**) corresponding % and counts (#, normalized to 5 x 10^5^ events) data for resident CD8^+^ T cells in WT vs. *Ikzf3^-/-^* infected lungs 1 d.p.i. (**e**) Frequency (%) and counts (#, normalized to 5 x 10^5^ events) for CD8^+^ T_VM_ cells in the lungs of WT and Aiolos-deficient mice 1 d.p.i. (**f**) Fold change in the number of CD8^+^ T_VM_ cells in the spleens and lungs of WT and Aiolos-deficient mice 1 d.p.i., relative to WT. Data representative of 5 independent experiments, n=14-16, mean ± SEM; unpaired Student’s t-test, **p≤0.01, ***p≤0.001 and ****p≤0.0001. (**g**) Schematic of murine influenza A virus infection and concurrent treatment with Fingolimod (FTY720). WT and *Ikzf3^-/-^*mice were intranasally infected with 100 PFU of PR8. Mice were intraperitoneally (i.p.) injected with FTY720 (1 mg/kg/day) or saline control daily starting at 1 day prior (day -1) to infection to two days (day 2) post-infection. (**h**) Data showing differences in the number (#, normalized to 3 x 10^5^ events) of resident CD8^+^ T cells in the lungs of WT and *Ikzf3^-/-^*mice in the control treated and FTY720 treated groups. Data shown for 2 independent experiments, n=6, mean ± SEM, Two-way ANOVA, Tukey’s multiple comparisons test, *p≤0.05, **p≤0.01.

Comparative analysis of T_VM_ cells in the spleens and lungs of WT and *Ikzf3^-/-^* mice revealed an average 2-fold increase in the number of T_VM_ cells in the spleens of Aiolos-deficient mice compared to WT. However, there was an approximate 12-fold average increase in the number of T_VM_ cells in the lungs of these mice compared to WT (Fig. 6f). To determine whether the increased lung T_VM_ population had trafficked from the spleen or expanded from the lung-resident population, we treated WT and *Ikzf3^-/-^* mice with Fingolimod (FTY720) or saline control via the intraperitoneal route daily starting one day prior to PR8 infection (Fig. 6g). FTY720 suppresses lymphocyte migration and has been widely used to study CD8^+^ T cell egress to infected tissues^44,45^. Following FTY720 administration, we infected mice with PR8 and examined the presence of resident CD8^+^ T cells in the lungs at 3 d.p.i. As before, infected animals treated with saline control displayed a significant increase in resident CD8^+^ T cells in the lungs of Aiolos-deficient mice relative to WT. In contrast, FTY720-treated animals exhibited diminished resident CD8^+^ T cell populations in the lungs of both WT and Aiolos-deficient mice, indicating that the T_VM_ population observed in the lungs migrated from secondary lymphoid organs in response to infection (Fig. 6h).

### Aiolos deficiency results in increased protection against influenza virus infection

To examine how increased T_VM_ populations in Aiolos-deficient settings may affect disease outcomes, we infected WT and Aiolos-deficient mice with PR8 (Fig. 7a). During infection, animal body weights were measured daily as an assessment of infection progression and severity. Consistent with past observations, Aiolos-deficient mice exhibited significantly less weight loss compared to WT mice as early as 5 d.p.i. (Fig. 7b)^26^. To determine whether the reduced weight loss was an outcome of reduced viral burden in the absence of Aiolos, we assessed viral load in the lung homogenates at 3 d.p.i. Indeed, both viral RNA (Fig. 7c) and infectious virus titers (Fig. 7d) were significantly reduced in Aiolos-deficient mice relative to WT. These findings demonstrate increased antiviral protection in the absence of Aiolos. As Aiolos expression is largely restricted to cells of the lymphoid lineage^46^, these early differences suggest an impact of Aiolos on an early-responding lymphocyte subset, such as T_VM_.

**Figure 7.**
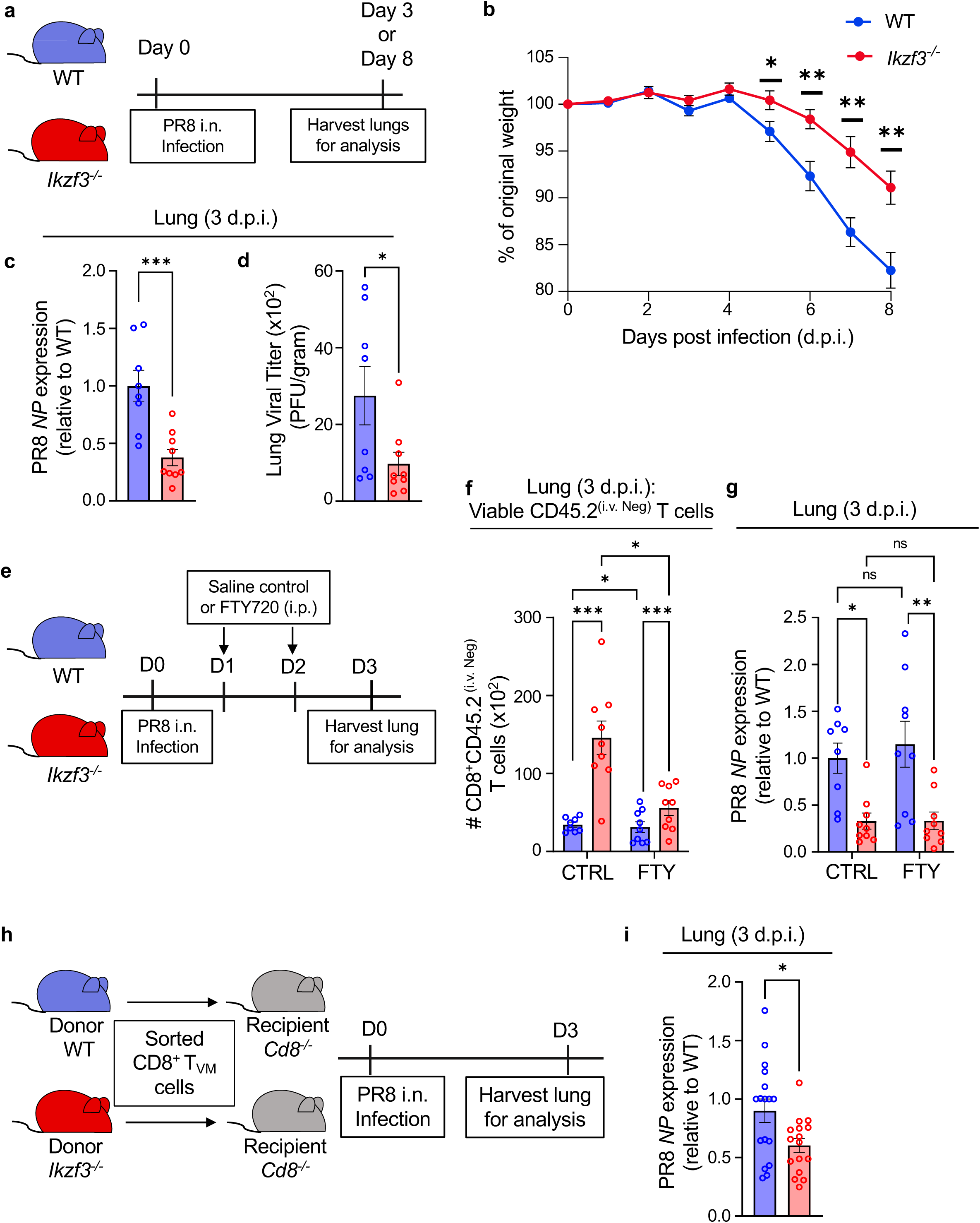
Aiolos-deficient CD8^+^ T_VM_ cells correlate with enhanced protection against influenza A virus infection. (**a**) Schematic of murine influenza A virus infection model. WT and Aiolos-deficient mice were infected with 30 PFU of PR8 intranasally and (**b**) weight of the mice was monitored daily up to 8 days post infection (d.p.i). Data representative of 4 independent experiments, n=14, mean ± SEM. *p≤0.05, **p≤0.01, multiple unpaired Student’s t-tests. (**c-d**) Determination of viral RNA and viral titer (PFU) at day 3 post-infection of WT and *Ikzf3^-/-^* mice with 100 PFU of PR8. (**c**) qRT-PCR analysis for influenza virus nucleoprotein (*NP*) on RNA isolated from the lung homogenates on day 3. Data normalized to *Rps18* and presented relative to WT. (**d**) Plaque-forming units (PFU) for viral titer in lung homogenates quantified 48 hours post-incubation with MDCK cells. Data represent the number of PFU per gram of lung tissue. Data shown for 2 independent experiments, n=8-9, mean ± SEM, Unpaired Student’s t-test, *p≤0.05, ***p≤0.001. (**e**) Schematic of murine influenza A virus infection and concurrent treatment with Fingolimod (FTY720). Mice were treated with saline or FTY720 (1 mg/kg/day) intraperitoneally (i.p.) at days 1 and 2 following influenza virus (100 PFU) infection. Tissues were collected after euthanasia at day 3 for flow cytometry and RNA analysis. (**f**) Data showing differences in the number (#, normalized to 3 x 10^5^ events) of CD45.2^(i.v. Neg)^ CD8^+^ T cells in the lungs of WT and *Ikzf3^-/-^* mice. (**g**) qRT-PCR analysis for influenza virus nucleoprotein (*NP*) expression on RNA isolated from the lung homogenates on day 3. Data normalized to *Rps18* and presented relative to WT. Data representative of 3 independent experiments, n=9, mean ± SEM; Two-way ANOVA, Tukey’s multiple comparisons test. *p≤0.05, **p≤0.01, ***p≤0.001. (**h**) Schematic of the adoptive transfer of sorted WT and *Ikzf3^-/-^*T_VM_ cells into recipient *Cd8^-/-^* mice and murine influenza virus infection for the determination of viral RNA levels as above. (**i**) qRT-PCR analysis for influenza virus nucleoprotein on RNA isolated from the lung homogenates on day 3 post-infection. Data normalized to *Rps18* and presented relative to WT. Data representative of 4 independent experiments, n=16-17, mean ± SEM; unpaired Student’s t-.test. *p≤0.05.

### Early lung-infiltrating Aiolos-deficient CD8^+^ T_VM_ cells correlate with enhanced protection against influenza virus infection

To determine whether the increased frequency of T_VM_ cells in the lungs of Aiolos-deficient mice (Fig. 6e) provides enhanced protection against viral infection, we infected WT and *Ikzf3^-/-^* mice with PR8 followed by FTY720 or saline administration at 1 d.p.i. This was done to limit our analysis to effects of the early-responding T_VM_ observed within the first day following infection as FTY720 treatment prevents later migration of lymphocytes to the site (Fig. 7e). Indeed, after treatment on days 1 and 2 post-infection, we observed increased resident CD8^+^ T cells in the lungs of Aiolos-deficient mice compared to WT in both saline and FTY720 treated groups (Fig. 7f). Given the limited presence of resident CD8^+^ T cells in the lungs of uninfected mice, the observed CD8^+^ T cell population likely represents infiltrating T_VM_ cells prior to FTY720 treatment. To assess the protective abilities of these early infiltrating CD8^+^ T cells, we determined viral RNA levels in the lungs of both the control and FTY720-treated mice. Similar to the control group, we observed significantly reduced viral RNA levels in the lungs of FTY720-treated Aiolos-deficient mice compared to WT (Fig. 7g). These findings suggest enhanced protection by Aiolos-deficient T_VM_ cells infiltrating the lungs as early as 1 d.p.i. Importantly, there was no difference in viral RNA levels between the control and FTY720-treated mice, demonstrating the efficacy of these early infiltrating T_VM_ cells.

To further assess the enhanced protective ability of Aiolos-deficient T_VM_ cells, we adoptively transferred WT or *Ikzf3^-/-^* CD8^+^ T_VM_ cells into recipient mice lacking CD8 (*Cd8^-/-^*) and infected recipients with PR8 1 day post-transfer (Fig. 7h). We then determined viral RNA levels in the lungs of the recipient mice at 3 d.p.i. We observed reduced viral RNA in the lung homogenates of mice that received Aiolos-deficient T_VM_ cells compared to mice that received WT T_VM_ cells (Fig. 7i), highlighting the superior ability of *Ikzf3^-/-^* T_VM_ to control virus titer in a cell-intrinsic manner. Together, these findings demonstrate a suppressive role for Aiolos in regulating CD8^+^ T_VM_ cell differentiation and function, particularly during early responses to viral infection.

## Discussion

Although Eomes and IL-15 signaling are well-recognized drivers of T_VM_ differentiation, the broader mechanisms governing CD8^+^ T_VM_ cell generation remain poorly understood. Here, we demonstrate that Aiolos functions as a negative regulator of T_VM_ cell generation and function. *In vivo*, Aiolos deficiency results in increased quantities of T_VM_ cells in the spleen under homeostatic conditions, and in both the spleen and lungs following influenza A virus infection. Importantly, the enhanced numbers of T_VM_ cells found in the lungs of Aiolos-deficient animals within 24 hours post-infection are not due solely to the expanded population in the spleen. In addition, Aiolos-deficient T_VM_ cells display enhanced effector function in response to cytokine stimulation. This combination of increased T_VM_ numbers and heightened effector function of T_VM_ cells correlates with improved viral clearance in Aiolos-deficient animals compared to WT controls at early stages of infection.

Mechanistically, our data supports a mechanism whereby Aiolos represses Eomes expression, thus limiting the formation of T_VM_ cells. Another consequence of this repression is the downregulation of the Eomes target gene *Il2rb* (encoding CD122/IL-15Rβ), leading to reduced IL-15/STAT5 signaling. Our data do not rule out the possibility that Aiolos may directly target *Il2rb* itself. Indeed, our prior studies in CD4^+^ T cells demonstrated that Aiolos is enriched at the *Il2ra* promoter, which correlated with reduced chromatin accessibility and transcription^26^. In another study, it was determined that Aiolos repressed IL-15 signaling and cytotoxic programming of intestinal intraepithelial lymphocytes, and that Aiolos binding site were proximal to that of STAT5^47^. Further, data from our lab also demonstrated a positive correlation between STAT5-binding sites and regulatory regions displaying increased chromatin accessibility in Aiolos-deficient CD4^+^ T cells^26^. Here we find the repression of STAT5 induced *Eomes* promoter response by Aiolos. Collectively, these findings support an antagonistic relationship between Aiolos and cytokine pathways that signal through STAT5.

While the necessity of Eomes in the promotion of T_VM_ cell differentiation is clear, it should be noted that Eomes harbors several other described roles in CD8^+^ T cell biology^14,35,36^. For example, while Eomes is generally considered a regulator of cytotoxic programming, it is also known to preferentially drive memory cell programming. This is in contrast to the related T-box factor T-bet, which promotes short-lived effector programming^48^. Within individual memory subsets, Eomes is known to promote circulating memory T cells while repressing tissue-resident cells^36,49–51^. As T_VM_ cells contribute to the formation of antigen-specific memory populations, particularly those that are lung-resident in the context of pulmonary infections, it remains to be seen whether Aiolos deficiency may impact the development of long-term memory CD8^+^ T cells^17^. To this end, CCR2^+^ T_VM_ cells have been reported to preferentially differentiate into T_RM_ cells following PR8 infection^17^. Our RNA-seq data revealed increased *Ccr2* transcript in Aiolos-deficient samples, thus supporting the possibility that loss of Aiolos may enhance T_VM_ differentiation into T_RM_ cells. However, given the role of Aiolos in repressing Eomes, it is intriguing to consider that in the absence of Aiolos, unchecked Eomes expression may inhibit the formation of a robust T_RM_ population, and perhaps, potentiate T_CM_ or T_EM_ subsets. Ultimately, future studies will be necessary to comprehensively assess the contribution of Aiolos to the generation of the full breadth of antigen-specific CD8^+^ T cell memory subsets.

T_VM_ cells have been implicated in various health conditions such as cancer and infections, where T_VM_ infiltration into the afflicted sites has been shown to improve the outcome of the disease^17–19,21^. In that regard, the findings of this study on the role of Aiolos opens a new avenue that could be utilized to enhance the therapeutic ability of these cells. In clinical settings, Aiolos, alongside Ikaros, is a known target of lenalidomide, a chemotherapeutic used for the treatment of multiple myeloma^52–54^. More recently, lenalidomide has been explored for lupus treatment and has demonstrated effectiveness in enhancing CAR T cell cytotoxicity in adoptive immunotherapy^55–57^. Our findings presented here suggest that targeting Aiolos may also influence the development and function of various CD8^+^ T cell subsets, with potential outcomes that could be either beneficial or detrimental, depending on the disease context. Beyond CD8^+^ T cells, Aiolos is expressed in other lymphoid populations, including NK cells and innate lymphoid cells (ILCs), which rely on IL-15 signaling and Eomes for their development and maintenance^54,58–64^. Thus, the effects of Aiolos may extend to these lymphoid populations. This study highlights the need for additional studies aimed at comprehensively understanding the contribution of Aiolos, and other Ikaros family members, in the differentiation and function of lymphocyte populations.

## Materials and Methods

### Mouse strains and cell lines

Wild-type CD45.2 C57BL/6J (JAX stock #000664) mice were originally obtained from the Jackson Laboratory. Aiolos-deficient mice were originally obtained from Riken BRC and backcrossed onto the CD45.2 C57BL/6J background for more than 10 generations to obtain CD45.2 Aiolos-deficient (*Ikzf3^-/-^*) mice. *Cd8^-/-^*mice (JAX stock #002665) were generously provided by Dr. Ginny Bumgardner and Dr. Hazem Ghoneim at The Ohio State University. All mice were housed in a specific-pathogen free (SPF) facility. Age- and sex-matched mice were used for both in vivo and in vitro studies to avoid any unintentional bias. All studies performed utilizing mice were done with the approval of the Institutional Animal Care and Use Committee (IACUC) at The Ohio State University in Columbus, OH. For the transfection studies, the EL4 murine T cell lymphoma line (TIB-39) was acquired from the American Type Culture Collection (ATCC) and maintained in complete RPMI (RPMI media [Thermo Fisher Scientific] containing 10% FBS [Life Technologies] and 1% penicillin/streptomycin [Life Technologies]). For transduction studies, the Platinum-E (Plat-E) Retroviral Packaging cell Line (RV-101, Cell Biolabs, Inc.) was maintained in complete DMEM (DMEM [Thermo Fisher Scientific], 10% FBS and 1% penicillin/streptomycin). MDCK (Madin–Darby canine kidney cells) cell line (ATCC, NBL-2) for viral titer determination was maintained in complete RPMI (RPMI, 10mM HEPES [Thermo Fisher Scientific], 1X L-glutamine [Thermo Fisher Scientific], 1X sodium pyruvate [Thermo Fisher Scientific], 1X MEM NEAA [Thermo Fisher Scientific], 10% FBS and 1% penicillin/streptomycin).

### CD8^+^ T cell isolation and culture

For overexpression studies, naive CD8^+^ T cells were isolated from the spleens and lymph nodes of 5-8-week-old *Ikzf3^-/-^*mice using the BioLegend MojoSort naive CD8^+^ T cell isolation kit according to the manufacturer’s recommendations. Naive CD8^+^ T cells (1.5 x 10^5^/mL) were plated in complete IMDM (IMDM [Life Technologies] with 10% FBS, 1% penicillin/streptomycin, and 0.05% (50 μM) 2-mercaptoethanol [Sigma-Aldrich]). Purified CD8^+^ T cells were plated in wells coated with anti-CD3 (5 μg/mL, clone 145-2C11, BD Biosciences) and anti-CD28 (2 μg/mL, clone 37.51, BD Biosciences) in the presence of IL-4-neutralizing antibody (5 μg/mL, clone 11B11, BioLegend) and IL-2-neutralizing antibody (5 μg/mL, JES6-1A12, BioLegend). IL-15 complex (IL-15C) was added to the indicated samples at a concentration of 100 ng/mL of IL-15. The IL-15C was generated as previously described^65^. Briefly, soluble recombinant IL-15 (447-ML, R&D) and recombinant mouse IL-15R⍺ Fc chimera (551-MR, R&D) were mixed at a 1:2 ratio and incubated at 37°C for 45 minutes immediately prior to use. Cells were cultured under the above conditions for 72 hours before downstream analyses.

For ex vivo functional assessment of CD8^+^ T_VM_ cells, bulk CD8^+^ T cells were isolated from the spleens of 5-8-week-old WT and *Ikzf3^-/-^* mice. Cells were purified using BioLegend MojoSort CD8^+^ T cell isolation kit according to the manufacturer’s recommendations. Bulk CD8^+^ T cells were treated with rmIL-12 (10 ng/mL, 419-ML, R&D), rmIL-18 (10 ng/mL, 9139-IL, R&D), and IL-15C (100 ng/mL) for 30 minutes to determine phosphorylation of STAT5 and STAT4 or for 16 hours to determine IFN-γ and granzyme B production. To preserve the phosphorylation of STAT proteins, cells were immediately fixed with 4% formaldehyde (37% formaldehyde, Fisher Scientific) post-stimulation and harvested for further analysis. For IFN-γ and granzyme B, cells were treated with protein transport inhibitors (PTI, 00-4980-93, Invitrogen) 4 hours prior to harvest for downstream analysis.

### Primary T cell Transduction

Aiolos overexpression experiments in primary CD8^+^ T cells were performed as previously described^26^. Briefly, pMIG-Aiolos was generated by cloning the Aiolos (*Ikzf3*) coding sequence into the pMSCV-IRES–GFP II (pMIG II, 52107, Addgene) vector backbone. The cloning of Aiolos was confirmed by Sanger sequencing and overexpression was validated via both qRT-PCR and immunoblot using anti-Aiolos antibody. Plat-E cells were used to package pMIG-Aiolos virus per the manufacturer’s instructions. Viral supernatants were collected 48 hours post-transfection and snap frozen. Viral supernatants were thawed immediately before use and added to murine CD8^+^ T cells, as detailed above, in the presence of polybrene (8 μg/mL, Sigma Aldrich). Transduction was performed at 24 and 48 hours after plating CD8^+^ T cells via spinfection for 2 hours at 800 x g at room temperature. At 48 hours, cells were treated with anti-IL-2 and IL-15C and harvested 24 hours after the second round of transduction for analysis.

### Promoter-reporter assay

To generate the Eomes promoter-reporter construct (pGL3-Eomes), the regulatory region of the *Eomes* gene (2 kb upstream of the transcriptional start site) was cloned into the Promega pGL3-Promoter vector (Promega). Expression vectors for Aiolos, Aiolos DNA-binding mutant (Aiolos^DBM^), and a constitutively active form of STAT5b (STAT5b^CA^) were generated as previously described^26,27^. Briefly, the coding sequence of Aiolos was cloned into the pcDNA3.1/V5-His-TOPO vector (Life Technologies). The coding sequence was confirmed by Sanger sequencing with T7 and BGH primers and then transferred into the pEF1/V5-His vector (Life Technologies). A similar technique was utilized for the generation of Aiolos^DBM^ and STAT5b^CA^ in conjunction with the Agilent QuikChange Site-Directed Mutagenesis kit (200519). For Aiolos^DBM^, point mutations were introduced into the first and second N-terminal zinc finger domains. For STAT5b^CA^, point mutations were introduced into STAT5b (H298R and S715F), resulting in constitutive phosphorylation of the Y699 residue. Using the Lonza 4D nucleofection system (program CM-120, buffer SF), EL4 T cells were nucleofected with the indicated vectors (V5-tagged Aiolos constructs, STAT5b^CA^, or empty vector control), along with pGL3-Eomes and a SV40-*Renilla* vector as a control for transfection efficiency. Following 22-24 hours of recovery, samples were harvested, and luciferase expression was analyzed using the Dual-Luciferase Reporter Assay System (Promega) according to the manufacturer’s instructions. Overexpression of the vectors was confirmed by Immunoblot analysis using an antibody against the V5 epitope tag (SV5-Pk1, Thermo Fisher Scientific) for Aiolos vectors and anti-STAT5b antibody (sc-1656, Santa Cruz Biotechnology) for STAT5b^CA^.

### RNA-Seq analysis

Bulk CD8^+^ T cells were isolated from the spleens of WT and *Ikzf3^-/-^* mice as described above. Cells were then stained for CD8α (PE, 1:300; clone 53-6.7; BioLegend), CD44 (V450; 1:300; clone IM7; BioLegend), CD62L (APCe780; 1:300; clone MEL-14; BioLegend), and CD49d (AF488; 1:100; clone R1-2; BD Biosciences) to identify T_VM_ populations, and were sorted using the Sony MA900 cell sorter for RNA extraction. For naive RNA samples, CD8^+^ T cells from the spleens of WT mice were isolated using the BioLegend MojoSort naive CD8^+^ T cell isolation kit as mentioned above. RNA was isolated from the sorted WT and Aiolos-deficient T_VM_ cells as well as WT naive CD8^+^ T cells using the Machery-Nagel Nucleospin RNA kit as per the manufacturer’s instructions. Purified RNA samples were submitted to Azenta Life Sciences for polyA selection, library preparation, and Illumina 2×150bp sequencing.

Data were pre-processed with Trimmomatic (0.36) and aligned with STAR (2.5.2b) to GRCm38. Transcript counts were generated with Subread (1.5.2) and quantified with DESeq2 (v 1.44.0) for analysis. Before analysis, expression data were filtered to exclude non-protein coding transcripts and lowly expressed genes (less than 100 counts). Genes with a DeSeq2 adjusted p<0.05 were considered differentially expressed. For gene set enrichment analysis (GSEA), differentially expressed genes were pre-ranked by multiplying the sign of the fold-change by −log10 (adjusted p value) and analyzed using the Broad Institute GSEA software (v 4.3.3) for comparison against “gene ontology,” and “immunologic signature” gene sets. Morpheus software (https://software.broadinstitute.org/morpheus) was used for heatmap generation and clustering by Euclidean distance using normalized log2 counts from DESeq2 analysis. Volcano plots were generated using −log10 (adjusted p value) and log2 fold change values from DESeq2 analysis in the VolcaNoseR software (https://huygens.science.uva.nl/VolcaNoseR/)^66^.

### Influenza virus infection and tissue preparation

Influenza A virus strain A/PR/8/34 (H1N1, termed “PR8”) was propagated in 10-day-old specific pathogen free embryonated chicken eggs (Charles River Laboratories) and titered on MDCK cells (BEI Resources, NIAID, NIH: Kidney [Canine], NR-2628). WT and Aiolos-deficient mice between 8-12 weeks of age were infected intranasally with 30 or 100 plaque-forming units (PFU) of PR8. For the adoptive transfer studies, WT and *Ikzf3^-/-^* T_VM_ cells were sorted as described above and retro-orbitally transferred into recipient *Cd8^-/-^*mice. The recipient mice were then infected via the intranasal route with 100 PFU of PR8. At designated times after infection, spleen and lung tissues were harvested as previously described^26^. To distinguish circulating from resident cells, mice were retro-orbitally injected with fluorophore-labeled anti-CD45.2 antibody (PE-anti CD45.2, 2 μg, clone #104, BioLegend) 3 minutes prior to euthanasia^17^. Tissues were then harvested and processed into single-cell suspensions. For spleen, single-cell suspensions were generated in tissue processing media (IMDM + 4% FBS) by passing the tissue through a 100 µm nylon mesh strainer followed by erythrocyte lysis in 0.84% NH_4_Cl. For lungs, whole lung tissue was incubated in HBSS (Gibco) with 1.3 mM EDTA followed by tissue dissociation in Collagenase IV supplemented media (RPMI + 4% FBS) using a gentleMACS Dissociator (Miltenyi Biotech) according to the manufacturer’s instructions. Samples were then filtered through a 40 μm nylon mesh strainer and layered with a Percoll density gradient to isolate lymphocytes. Cells were centrifuged at 500 x g for 20 minutes with brakes off to allow for separation of mononuclear cells. Cells were then collected, and erythrocyte lysis was performed as previously described. Cells were resuspended in FACS buffer (PBS + 4% FBS) prior to staining for flow cytometry. Spleens from uninfected mice were processed similarly as above and single-cell suspensions were resuspended in FACS buffer for flow cytometry analysis.

### FTY720 administration

Mice were injected with saline control or Fingolimod (FTY720, Selleckchem) at 1 mg/kg in saline daily via the intraperitoneal route at indicated time points. Tissues were then collected post-euthanasia at indicated times for processing and analysis.

### Flow cytometry

For flow cytometry analysis of lung and spleen samples, cells were first incubated with anti-mouse CD16/32 Fc block (clone 93; 101320; BioLegend) for 5 minutes at 4°C. Cells were then incubated for 30 minutes at 4°C for extracellular staining with following antibodies: anti-CD8α (BV570, 1:300; clone 53-6.7; BioLegend), anti-CD3 (BV605; 1:100; clone 17A2; BioLegend), anti-CD44 (BV785; 1:300; clone IM7; BioLegend), anti-CD62L (BV650; 1:300; clone MEL-14; BioLegend), anti-CD49d (BUV395; 1:100; clone R1-2; BD Biosciences), anti-CD122 (FITC; 1:00; clone 5H4; BD Biosciences), and Ghost viability dye (Red 780; 1:780; Tonbo Biosciences). Cells were then washed twice with FACS buffer prior to intracellular staining. For intracellular staining, cells were fixed and permeabilized using the eBioscience Foxp3 transcription factor staining kit (Thermo Fisher Scientific) for 30 minutes or overnight at 4°C. After fixation, samples were stained with the following antibodies in 1X permeabilization buffer (Thermo Fisher Scientific) for 30 minutes at room temperature: anti-Aiolos (AF647; 1:100; clone S48-791; BD Biosciences), anti-Eomes (PE-Cy7; 1:100; clone DAN11MAG; Thermo Fisher Scientific), anti-IFN-γ (APC; 1:500; clone XMG1.2; BioLegend) and anti-granzyme B (PE; 1:250; clone GB11; Thermo Fisher Scientific). Cells were then washed with 1X permeabilization buffer and FACS buffer once before resuspending in FACS buffer for analysis. To determine the phosphorylation levels of STAT proteins, cells were immediately fixed with 4% formaldehyde (37% formaldehyde, Fisher Scientific) post-stimulation to preserve the phosphorylation of STAT proteins. Cells were then harvested and washed with 1X PBS prior to incubation with anti-CD49d and anti-CD44 antibodies for 30 minutes at 4°C. Following the extracellular staining, cells were permeabilized with cold methanol overnight at - 20°C. These cells were then incubated with antibodies against pY-STAT5 (Y694, APC, clone: SRBCZX, Thermo Fisher Scientific), pY-STAT4 (PY933, PerCP-Cy5.5, clone: 38/p-Stat4, BD Biosciences), and CD8α for 30 minutes before resuspending in FACS buffer for analysis. Samples were run on a BD FACS Canto II or BD FACS Symphony and analyzed using FlowJo software (version 10.8.1).

### Virus titers and Viral RNA determination

Virus titers and virus RNA were determined as previously described^67–69^. Briefly, mice were euthanized at designated time points post-infection and lungs were harvested in 1 mL PBS + 0.5% BSA in Precellys tubes (Bertin Corp) for homogenization using the Precellys homogenizer. The lung supernatants were collected, snap frozen, and stored at -80°C prior to use. For the determination of virus titers, MDCK cells were plated at a density of 7 x 10^5^ cells/well in a 6-well plate. Virus supernatants were 10-fold serially diluted and added to MDCK cells in the presence of infection media (1X PBS, 35% BSA [VWR], 1X Ca/Mg). Following 1 hour incubation at 37°C the viral supernatant was removed, and cells were washed with 1 X PBS. An overlay (2% Oxoid agar [Thermo Fisher Scientific], 2X MEM [Lonza], TPCK Trypsin [Thermo Fisher Scientific], DEAE-Dextran [Thermo Fisher Scientific], 5% sodium bicarbonate and H_2_0) was added to the wells. After 48 hours of incubation, the overlay was removed, and cells were fixed with 3% paraformaldehyde (PFA). Cells were blocked with 5% non-fat milk in 1X PBS and incubated with chicken anti-PR8 antibody (BEI Resources, NR-3098) and rabbit anti-chicken peroxidase (Jackson ImmunoResearch, 303-035-003). The number of plaques formed were quantified following the addition of TrueBlue Peroxidase (Thermo Fisher Scientific).

For viral RNA determination, lungs were harvested in 1 mL Trizol (Life Technologies) and processed as above. 1-bromo-3-chloropropane (Sigma Aldrich) was added to the clarified virus supernatant. The aqueous layer was collected post-centrifugation, ethanol was added, and the mixture was transferred onto a spin column from the Machery-Nagel Nucleospin RNA kit. RNA was extracted according to the manufacturer’s instructions. cDNA was prepared and qRT-PCR analysis was then performed to determine virus RNA levels.

### RNA isolation and Quantitative Real-Time PCR (qRT-PCR)

RNA was isolated as described above utilizing the Machery-Nagel Nucleospin RNA kit per the manufacturer’s guidelines. cDNA was then generated using the Superscript IV First Strand Synthesis System (Thermo Fisher Scientific). SYBR Select Mastermix for CFX (ThermoFisher) was used for qRT-PCR reactions with 10 ng cDNA per reaction on the CFX Connect (BioRad). The following primers were used: (IAV *NP* primers-Forward: 5’ – GATTGGTGGAATTGGACGAT – 3’, Reverse: 5’ – AGAGCACCATTCTCTCTATT – 3’, *Rps18* primers-Forward: 5’ – GGAGAACTCACGGAGGATGAG –3’, Reverse: 5’ – CGCAGCTTGTTGTCTAGACCG -3’). Outputs were normalized to *Rps18* and are presented as relative to their respective control, as indicated.

### Immunoblot analysis

For immunoblot analysis, cells were harvested, lysed in 1X SDS loading dye, (50 mM Tris [pH 6.8], 100 mM DTT, 2% SDS, 0.1% bromophenol blue, 10% glycerol) and boiled for 15 minutes. Lysates were separated via SDS-PAGE on 10% Bis-Tris Bolt gels (Thermo Fisher Scientific) and then transferred onto a 0.45 µm nitrocellulose membrane. Membranes were blocked with 2% nonfat dry milk in 1X TBST (10 mM Tris [pH 8.0], 150 mM NaCl, 0.05% Tween-20). Membranes were then incubated with the following antibodies to detect proteins: α-V5 (1:20,000; clone SV5-Pk1, Invitrogen), α-STAT5b antibody (1:10,000; sc-1656, Santa Cruz Biotechnology), goat ⍺-mouse HRP (1:10,000; 115-035-174, Jackson Immunoresearch), ⍺-β-actin-HRP (1:15,000; clone: 2D1D10, GenScript). Clarity Max Western ECL substrate (BioRad) was used for the visualization of protein bands.

### Statistics and reproducibility

All statistical analyses were performed using the GraphPad Prism software (version 9.5.1). Unpaired Student’s t-test was performed for single comparisons. For multiple comparisons, one-way or two-way ANOVA with Tukey’s multiple comparisons test was performed, as appropriate. Error bars indicate the standard error of the mean. P values ≤0.05 were considered statistically significant.

### Software summary

Data were collected utilizing the following software programs: BD FACSDiva (version 8.0.2), BioRad Image Lab (version 6.0.1, build 34), BioRad CFX Manager (version 3.1). Analyses and/or manuscript preparation were conducted using Microsoft Office Suite (including Microsoft Word, Microsoft Excel, and Microsoft PowerPoint), BD FlowJo (version 10.8.1), Integrative Genomics Viewer (version 2.18.2), Morpheus (Broad Institute), Gene Set Enrichment Analysis (GSEA, v4.3.3; Broad Institute) and VolcanoseR (https://huygens.science.uva.nl/VolcaNoseR/)^66^. All statistical analyses were performed using the GraphPad Prism software (version 10.3.1). Data preparation for this manuscript did not require the use of custom code or software.

## Data Availability

Data are available in the article or at a reasonable request to the corresponding author. RNA-seq data has been deposited in the GEO repository under accession number GSE284441. The following publicly available ChIP-seq data were used in this study: Aiolos ChIP-Seq: GSM5106065^37^ and STAT5b ChIP-Seq: GSM7887512^39^.

## Acknowledgements

The authors would like to thank all members of the Oestreich Laboratory, as well as colleagues in the Department of Microbial Infection and Immunity, for critical feedback. The authors would also like to thank members of the Yount Laboratory for help with influenza virus preparation and members of the Hemann Laboratory (Abigail Solstad) and Forero Laboratory (Matthew McFadden) for their technical advice on influenza virus titer and RNA determination. The authors would also like to thank Dr. Ginny Bumgardner and Dr. Hazem Ghoneim for providing *Cd8^-/-^* mice. This work was supported by grants from the National Institutes of Health (R01AI134972) and funds through The Ohio State University College of Medicine and The Ohio State University Comprehensive Cancer Center to K.J. Oestreich. S. Pokhrel was supported by funding through the American Heart Association Postdoctoral Fellowship (https://doi.org/10.58275/AHA.24POST1196412.pc.gr.190821) and The Ohio State University Infectious Disease institute (IDI) Interdisciplinary Research Seed Grant. S. Pokhrel, M.R. Leonard, and J.A. Tuazon were supported by the National Institutes of Health T32: AI165391, Interdisciplinary Program in Microbe-Host Biology. J.A. Tuazon was also supported by National Institutes of Health F30: AI172189-01A1 and The Ohio State University Susan Huntington Dean’s Distinguished University Fellowship. G. Dileepan was supported by The Ohio State University Dean’s Distinguished University Fellowship.

## Author Contributions

S. Pokhrel assisted with the design of the study, performed experiments, analyzed data, and wrote the manuscript. D.M. Jones, M.R. Leonard, J.A. Tuazon, K.A. Read, G. Dileepan, R.T. Warren, Q. Gong performed experiments and analyzed data. J.S. Yount, G. Xin, A. Forero, H.E. Ghoneim, E.A. Hemann assisted with the influenza virus infection experiments, data analysis, and reviewed the manuscript. P.L. Collins assisted with the RNA-Seq data analysis and reviewed the manuscript. K.J. Oestreich supervised the research, designed the study, analyzed data, and wrote the manuscript.

