## Supplementary Figures for "Aiolos restricts the generation of antigen-inexperienced, virtual memory CD8^+^ T cells"

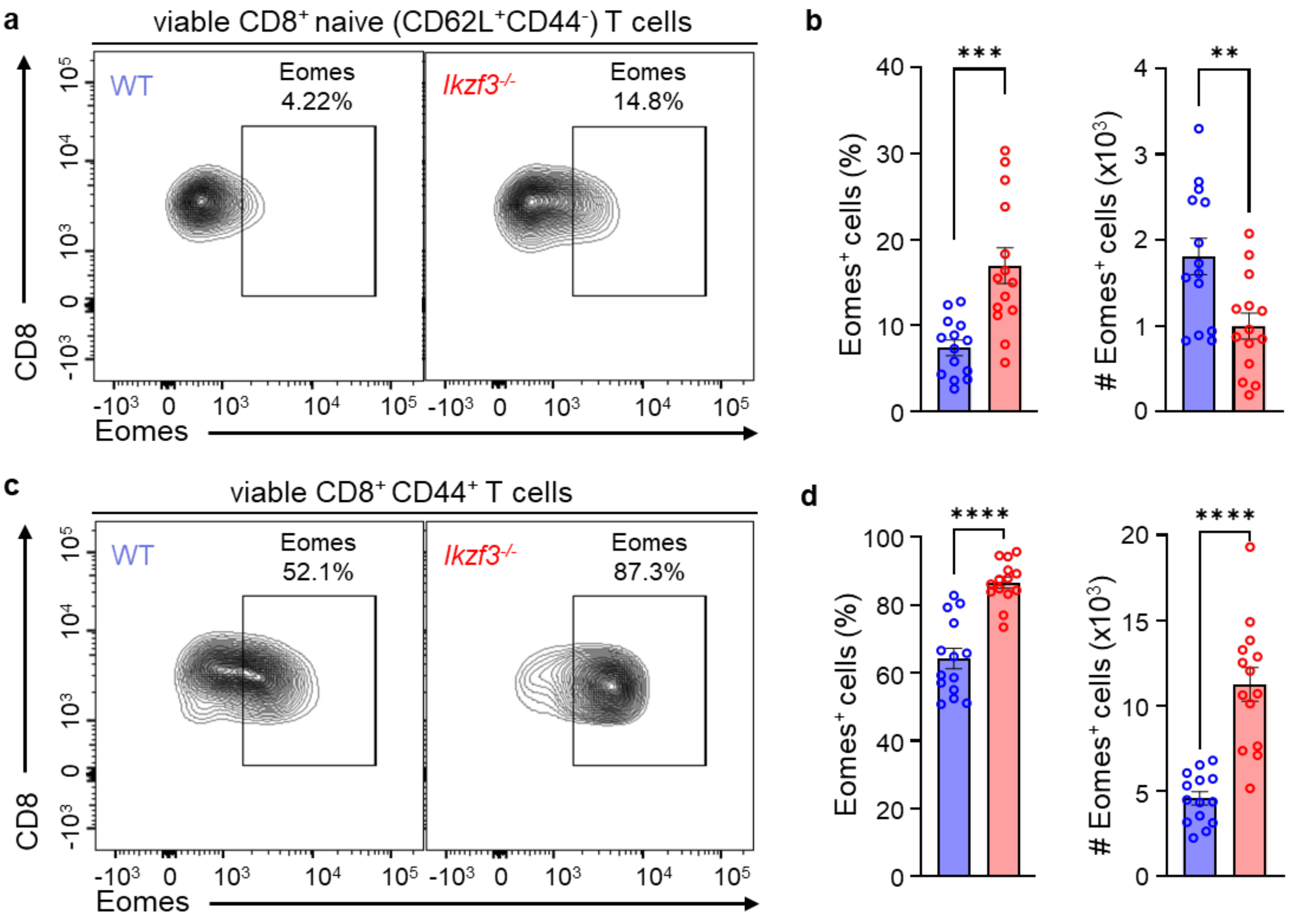


**Supplementary figure 1: Eomes and Aiolos expression inversely correlate in CD8$^+$ T cells.** Flow cytometry analysis of Eomes expression in CD8$^+$ naive (CD62L$^+$CD44$^-$) and activated (CD44$^+$) T cells from the spleens of uninfected WT and *Ikzf3*$^{-/-}$ mice. Representative flow cytometry contour plots and corresponding frequency and numbers (#, normalized to 6 x $10^5$ events) of Eomes-expressing (**a-b**) naive and (**c-d**) effector CD8$^+$ T cells. Data shown for 4 independent experiments, n=14, mean ± SEM, unpaired Student's t-test, **p≤0.01, ***p≤0.001, ****p≤0.0001.

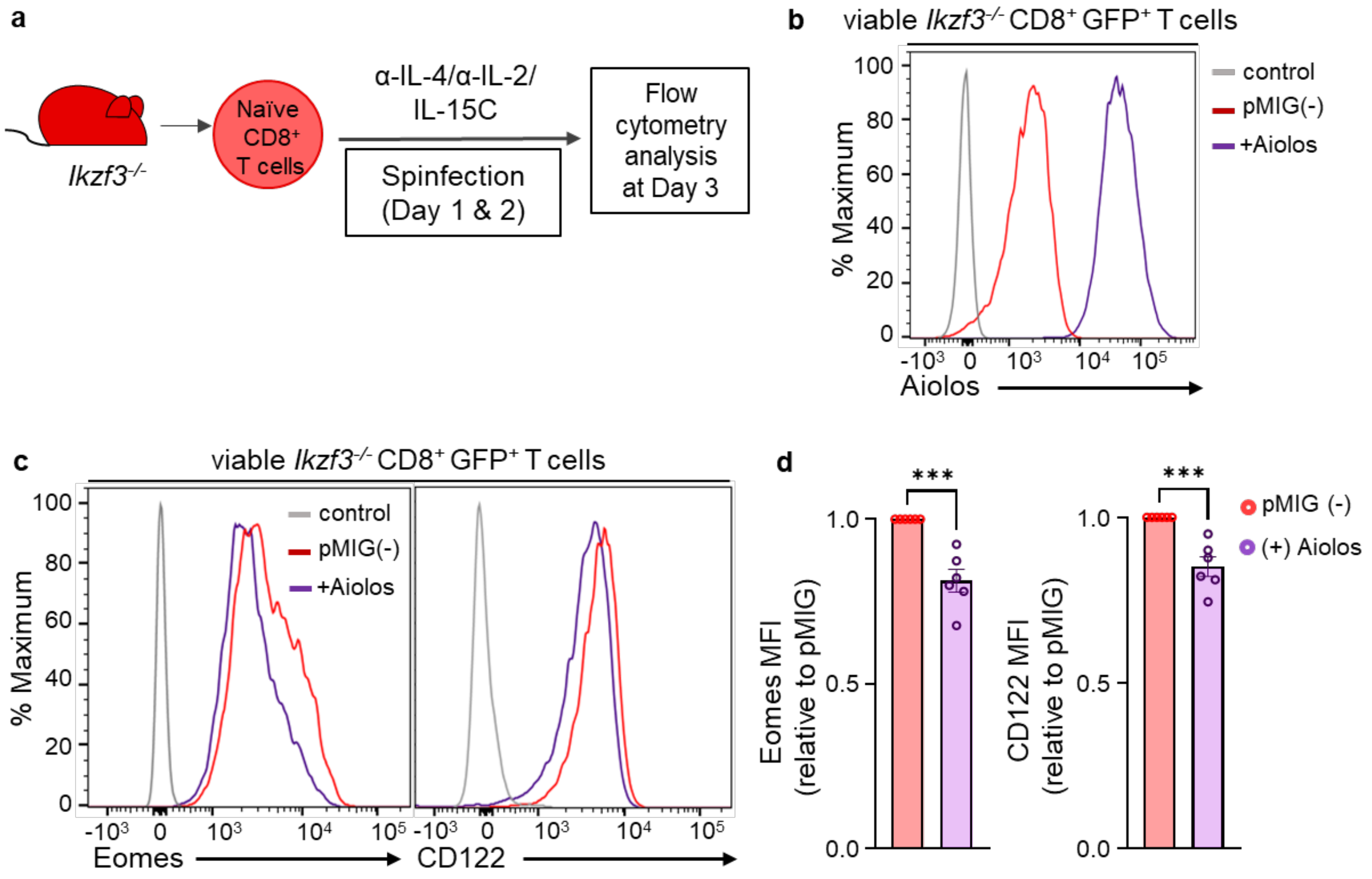


**Supplementary figure 2: Aiolos overexpression results in decreased Eomes and CD122 expression in Aiolos-deficient CD8$^+$ T cells.**

(**a**) Schematic of overexpression of Aiolos in Aiolos-deficient CD8$^+$ T cells. Naive CD8$^+$ T cells isolated from the spleens of *Ikzf3$^{-/-}$* mice were spinfected with viral supernatants for Aiolos overexpression. (**b**) Representative histogram overlay confirming Aiolos overexpression in Aiolos-deficient CD8$^+$ T cells. (**c**) Histogram overlay for Eomes and CD122 and (**d**) associated MFI fold change data following the overexpression of Aiolos in Aiolos-deficient CD8$^+$ T cells *in vitro*. Data shown for 6 independent experiments, n=6, mean ± SEM, unpaired Student's t-test, ***p≤0.001.

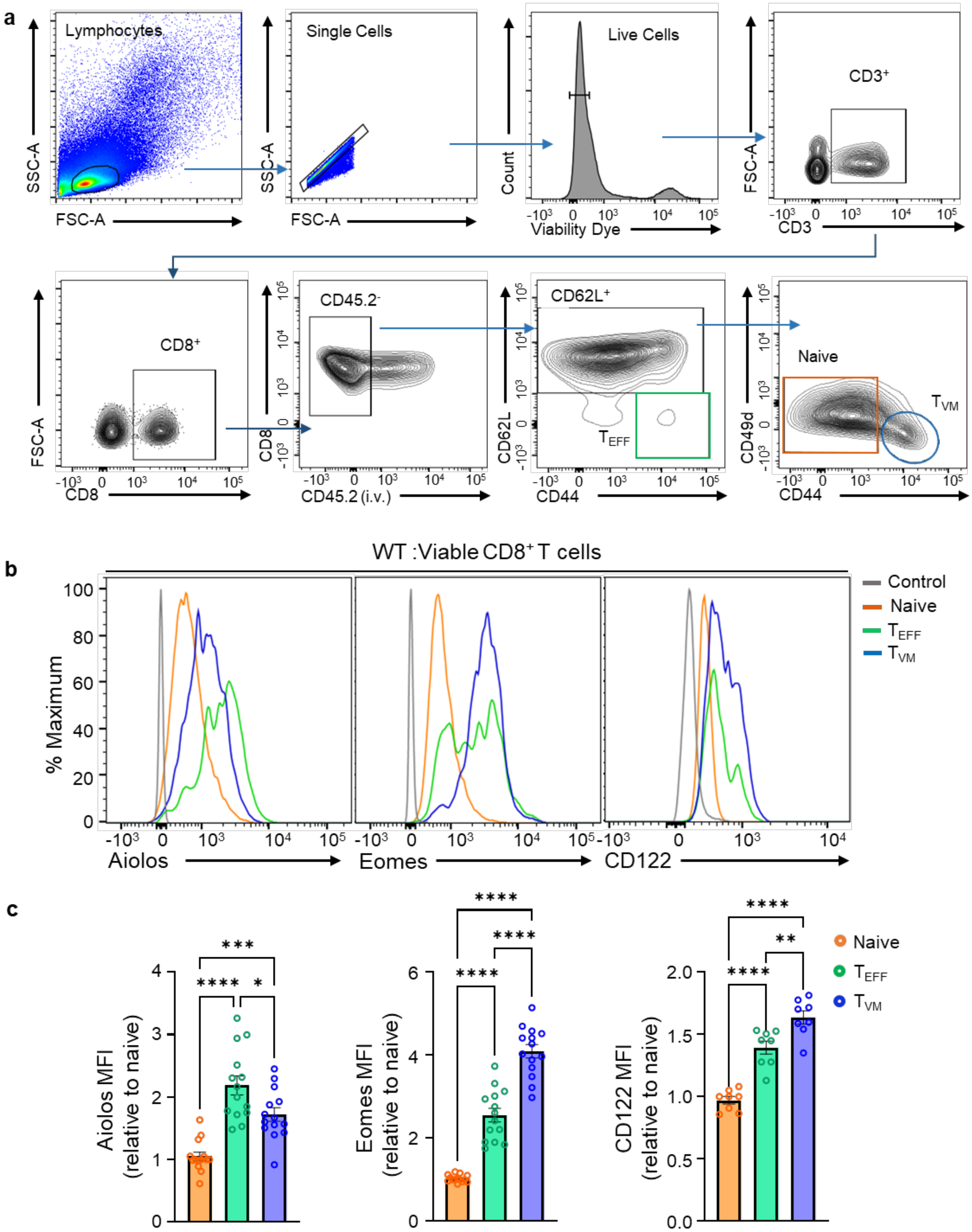
a
Lymphocytes
SSC-A
FSC-A
Single Cells
SSC-A
FSC-A
Live Cells
Count
Viability Dye
CD3+
FSC-A
CD3
CD8+
FSC-A
CD8
CD45.2-
CD8
CD45.2 (i.v.)
CD62L+
CD62L
TEFF
CD44
Naive
TVM
CD49d
CD44
b
WT :Viable CD8+ T cells
% Maximum
Aiolos
Eomes
CD122
Control
Naive
TEFF
TVM
c
Aiolos MFI
(relative to naive)
Eomes MFI
(relative to naive)
CD122 MFI
(relative to naive)
Naive
TEFF
TVM

**Supplementary Figure 3: Representative gating strategy and expression of Aiolos, Eomes, and CD122 in different CD8$^+$ T cell subsets.**

(**a**) Gating strategy for CD8$^+$ T cells from uninfected or infected WT and *Ikzf3*$^{-/-}$ mice. Mice were retro-orbitally injected with PE-labeled anti-CD45.2 antibody prior to euthanasia to distinguish circulating (CD45.2$^{(i.v.\ Pos)}$) cells from resident (CD45.2$^{(i.v.\ Neg)}$) cells. CD8$^+$ naive, effector ($T_{EFF}$) and virtual memory ($T_{VM}$) cells were distinguished using CD62L, CD44, and CD49d markers as: Naive (CD62L$^+$CD44$^-$CD49d$^-$), effector (CD62L$^-$CD44$^+$), and $T_{VM}$ (CD62L$^+$CD44$^+$CD49d$^{Lo}$). (**b-c**) Comparison of expression of Aiolos, Eomes, and CD122 across different CD8$^+$ T cell subsets in the spleens of uninfected WT mice. (**b**) Representative histogram overlays and (**c**) corresponding MFI fold changes for indicated proteins on naive, $T_{EFF}$, and $T_{VM}$ cells. Data shown for 2-4 independent experiments, n=14 (for Aiolos and Eomes) and n=8 (for CD122), mean ± SEM; One-way ANOVA, Tukey's multiple comparisons test, *p≤0.05, **p≤0.01, ***p≤0.001 and ****p≤0.0001.

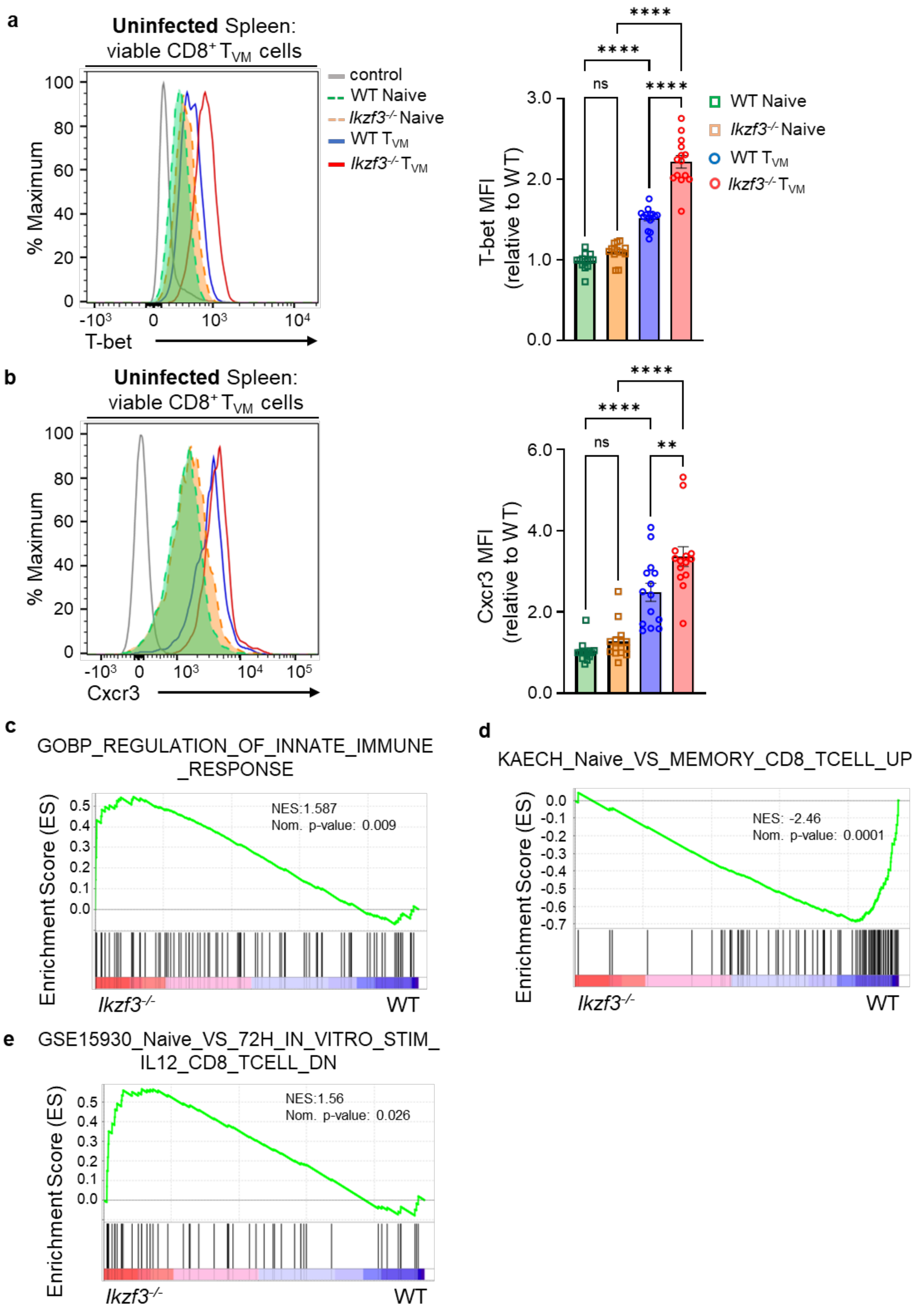
a
Uninfected Spleen:
viable CD8+ TVM cells
control
WT Naive
Ikzf3-/- Naive
WT TVM
Ikzf3-/- TVM
% Maximum
T-bet
T-bet MFI (relative to WT)
ns
****
**
b
Uninfected Spleen:
viable CD8+ TVM cells
Cxcr3
Cxcr3 MFI (relative to WT)
c
GOBP_REGULATION_OF_INNATE_IMMUNE_RESPONSE
NES:1.587
Nom. p-value: 0.009
Enrichment Score (ES)
Ikzf3-/-
WT
d
KAECH_Naive_VS_MEMORY_CD8_TCELL_UP
NES: -2.46
Nom. p-value: 0.0001
e
GSE15930_Naive_VS_72H_IN_VITRO_STIM_IL12_CD8_TCELL_DN
NES:1.56
Nom. p-value: 0.026

**Supplementary Figure 4: Aiolos suppresses T-bet and CXCR3 expression in $T_{VM}$ cells and regulates various gene pathways.**

(**a-b**) Representative histogram overlays and associated median fluorescence intensity (MFI) fold change for (**a**) T-bet and (**b**) CXCR3 expression in the indicated cell types, relative to WT naive $CD8^+$ T cells. Data shown for 4 independent experiments, n=14, mean ± SEM; two-way ANOVA with Tukey's multiple comparisons test, **$p \leq 0.01$, ****$p \leq 0.0001$. (**c-e**) Enrichment plots for indicated gene sets generated using the GSEA software for comparison against 'gene ontology', and 'immunological signature' gene sets of the pre-ranked genes.

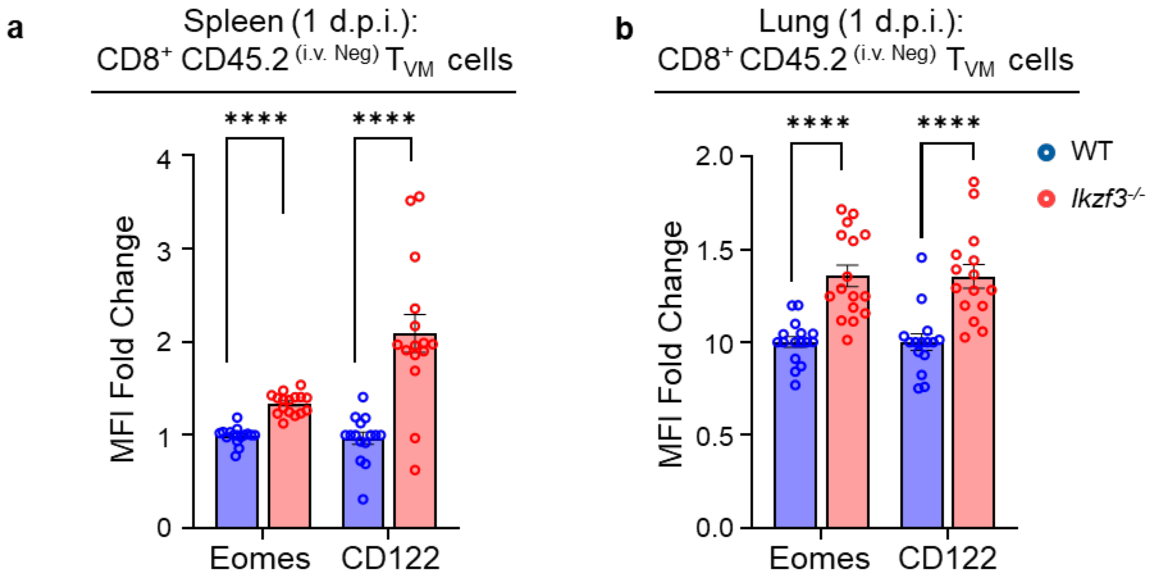


**Supplementary Figure 5: Aiolos suppresses the expression of key $T_{VM}$ regulatory factors.**

Median fluorescence intensity (MFI) fold change for Eomes and CD122 expression relative to WT in $CD8^+$ $T_{VM}$ cells from (**a**) the spleen and (**b**) lungs of WT vs. Aiolos-deficient mice 1 day following influenza virus challenge. Data representative of 5 independent experiments, n=14-16, mean ± SEM; unpaired Student's t-test, ****$p \leq 0.0001$.
